# Coordinated immune-epithelial dynamics in the nasal epithelium protect against respiratory virus infection

**DOI:** 10.64898/2026.06.08.726938

**Authors:** Yao Yu Yeo, Hsin-Yi Chen, Bokai Zhu, Yang Wang, Ivan T Lee, Tsuguhisa Nakayama, Yuchen Wang, Huaying Qiu, Kate B Juergens, Sei Chung, Carol H Yan, Clara S Kummerer, Hendrik A Michel, Andrew Z Ma, Wenrui Wu, Jayakar V Nayak, Garry P Nolan, David R McIlwain, Alexandar Tzankov, Matthias S Matter, Chien-Ting Wu, Sizun Jiang

## Abstract

Epithelial remodeling is a hallmark of host antiviral responses that strengthen barrier defenses against infection. Our current understanding of viral pathophysiology within the nasal epithelium, the initial site of most respiratory viral infections, remains incomplete due to limited understanding of the native tissue architecture and protective epithelial remodeling during immune events. Leveraging coronavirus disease 2019 (COVID-19) as a model, we applied spatial multi-omics on nasal cross-sectional tissues to characterize the immune–epithelial landscape during infection, identifying a coordinated increase in goblet cells and suppressive macrophages associated with elevated IL13 in the immune compartment. Our results further reveal that goblet cell and suppressive macrophage enrichment are spatially linked in proximity to IL13–expressing CD4 T cells, consistent with IL13-driven remodeling *in situ*. Using a primary human nasal air–liquid interface model, we demonstrate that IL13 alone is sufficient to remodel epithelial composition and morphology, subsequently restricting viral infection by reshaping the apical mucus barrier of the nasal epithelium. Our findings uncover a spatially organized, IL13–driven circuit for immune-epithelial remodeling as a protective barrier against respiratory viral infections.

## Introduction

A fundamental aspect of airway health vs. disease is maintenance of epithelial integrity against invading pathogens coordinated by the immune system. Immune-epithelial remodeling is recognized to underpin tissue regeneration and diseases including chronic rhinosinusitis, asthma, and COVID-19 (1–4). These interactions can involve epithelial secretion of soluble paracrine signals that shape immune infiltration to preserve tissue homeostasis or drive chronic inflammation. During acute respiratory viral infection, the epithelial layer undergoes dynamic remodeling that includes robust interferon responses accompanied by a rapid recruitment of inflammatory lymphoid and myeloid populations to promote antiviral defense (5, 6). Despite the growing recognition of immune-epithelial communications in shaping host responses to respiratory viruses, the mechanisms that link immune signaling to epithelial remodeling during infection remain incompletely understood, in part due to the inherent complexity of dissecting these interactions *in situ*, and technical limitations to resolve these processes within the native tissue context.

The nasal epithelium represents a major site of infection and early host–pathogen interaction across respiratory viruses (7) despite being a specialized barrier against pathogen invasion, making it a critical interface for immune coordination and epithelial remodeling. Amongst the human respiratory viruses, severe acute respiratory syndrome coronavirus 2 (SARS-CoV-2) provides a relevant context to dissect these critical processes for two key reasons. Firstly, the nasal epithelium is the site of primary infection where localized robust immune responses are first observed (8–11), and secondly, it is the causative agent of COVID-19, which has had transformative effects globally since its emergence in 2019 (12–14). Despite a strong tropism for nasal tissues, much of the focus on COVID-19 pathophysiology, as with other respiratory viruses, has centered on the lungs and bronchi, where infection can lead to acute respiratory distress and extensive pulmonary damage (4). Beyond localized tissue injury, COVID-19 can also induce cytokine storms, acute lesions, and systemic inflammation (4, 15, 16), leading to extensive tissue disruption that makes the relatively confined and fragile nasal epithelium a technically challenging tissue site for study. Prior investigations on nasal airways have also largely relied on nasopharyngeal swabs or nasal washes, which mainly sample the air surface liquid and superficial epithelium. These approaches fail to capture the underlying complexities of the intact tissue microenvironment and immune-epithelial interactions, thus limiting our understanding of how the nasal epithelium responds to respiratory viral infections.

To address these challenges and assess how acute viral infection reshapes the nasal epithelium, we first screened post-mortem nasal tissues from individuals who succumbed during the early wave of the COVID-19 pandemic, and carefully selected those with preserved morphology (17). We next applied spatial multi-omics using the GeoMx spatial transcriptomics (18) and CODEX spatial proteomics (19) platforms to examine immune-epithelial interactions and tissue remodeling within the native diseased tissue, followed by mechanistic dissection using a primary human nasal epithelium air-liquid interface culture system. Our findings reveal a spatially organized IL13–driven immune–epithelial circuit *in situ* that reshapes nasal epithelial composition to induce mucus secretion and establish a protective barrier against respiratory viral infection. Our integrative approach highlights spatial immune–epithelial interactions as a key axis shaping antiviral barrier function, thus advancing a framework towards understanding the native tissue-level host responses against respiratory viral infections in the nasal airway.

## Results

### Spatial transcriptomics capture respiratory viral-linked nasal immune-epithelial dynamics

To dissect epithelial and immune functional remodeling upon respiratory viral infection, we first applied the GeoMx spatial transcriptomics platform (18) on a cohort of formalin-fixed paraffin-embedded (FFPE) nasal epithelium samples from 10 COVID-positive and 4 COVID-negative individuals **(Fig. 1A)**. As the nasal epithelium is the initial site of SARS-CoV-2 infection with pervasive tissue damage and cell death (8, 9), we applied a probe panel targeting 1,800 genes with five probes per target gene to maximize RNA capture quality. Tissues were additionally stained with antibodies against pan-cytokeratin (panCK) and CD45 to visualize epithelial and immune compartments respectively for compartment-specific transcriptome capture.

**Figure 1.**
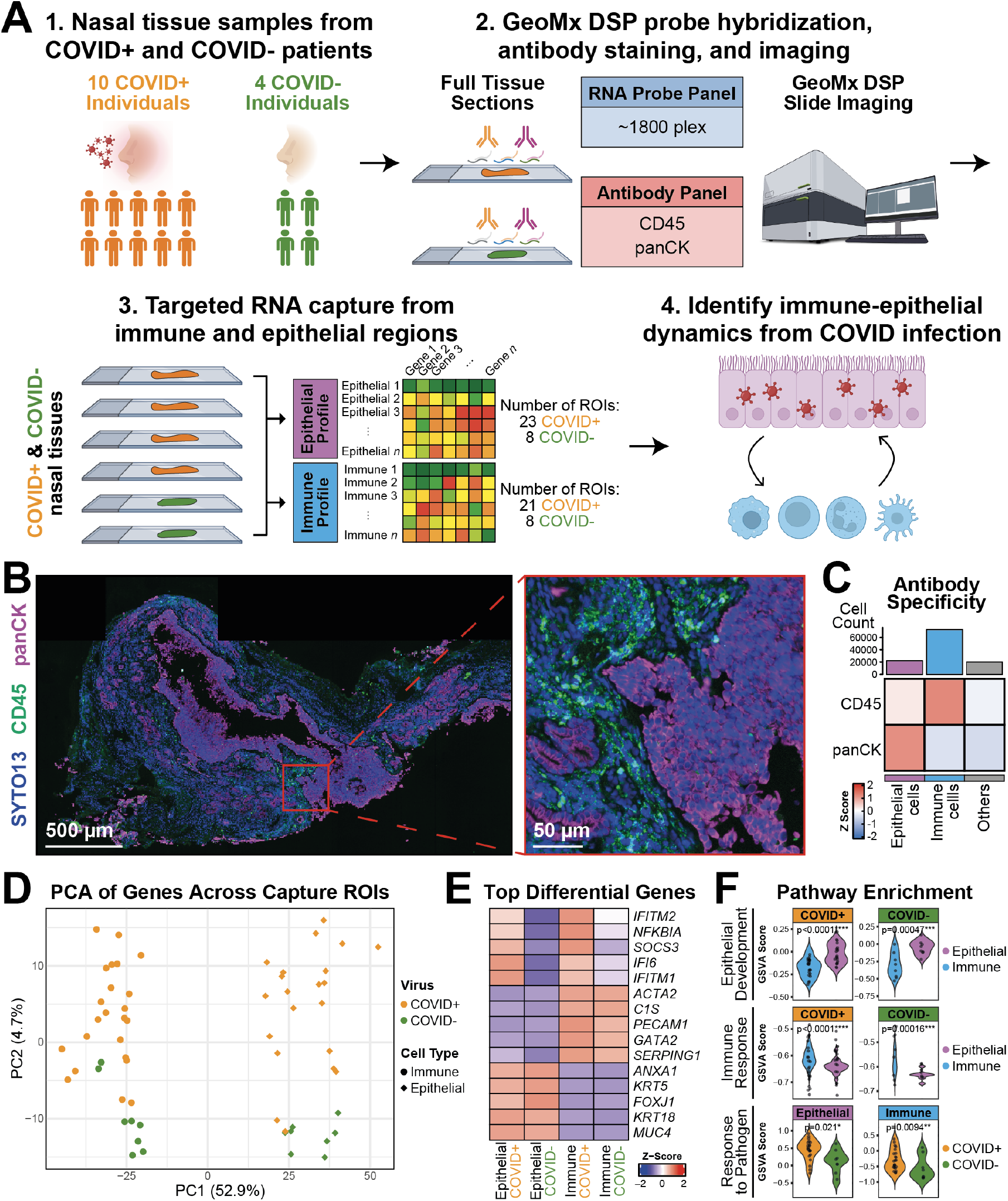
Application of GeoMx spatial transcriptomics to identify differences between COVID-positive and COVID-negative nasal tissues. **(A)** Strategy to spatially dissect the COVID-positive and COVID-negative nasal epithelium. (1) Biopsies from a cohort of COVID-positive (n=10) and COVID-negative (n=4) patients were (2) stained with a 1800-plex RNA probe panel and two morphology antibodies to enable (3) acquisition of the epithelial and immune transcriptomes using the GeoMx, where upon (4) computational analysis revealed key differences between COVID-positive and COVID-negative tissues. **(B)** Representative GeoMx immunofluorescence image of the nasal epithelium, with markers for nuclei (SYTO13), immune cells (CD45), and epithelial cells (panCK) shown. **(C)** Relative z-score expression levels of antibody markers for the annotated cell phenotypes in this GeoMx dataset. **(D)** PCA plot showing separation by tissue type along PC1 and viral status along PC2. **(E)** Heatmap of top 5 differentially expressed genes stratified by immune, epithelial, and viral status. **(F)** Comparison of representative Gene Ontology pathways stratified by tissue type or viral status. A one-sided Wilcoxon rank sum test with Benjamini-Hochberg correction was conducted for each comparison (* *p ≤* 0.05, ** *p ≤* 0.01, *** *p ≤* 0.001, **** *p ≤* 0.0001).

We assessed the specificity of spatial transcriptome capture on multiple levels. The antibody staining specificity was first confirmed visually **(Fig. 1B)** and consistent with the expected enrichment of antibody markers for epithelial and immune compartments **(Fig. 1C)**. Principal component analysis (PCA) was then applied to the 60 capture regions, revealing clear separation of transcriptomes by cell type across PC1 and viral status across PC2 **(Fig. 1D)**. Analysis of the top differentially expressed genes in each cell population showed an expected enrichment of genes that stratify viral status and cell population, including interferon-induced genes *(IFITM1, IFITM2, IFI6)* across COVID-positive samples, the complement system *(C1S)* across immune regions, and keratins *(KRT4, KRT5)* across epithelial regions **(Fig. 1E)**. Evaluation of representative gene signatures for epithelial, immune, and viral status (20) also demonstrated significant differences concordant to their respective capture regions **(Fig. 1F)**. The stringent validation of transcriptome capture quality in these technically challenging tissues demonstrate the robustness of this approach and data for further dissection of epithelial and immune dynamics in viral-infected, post mortem nasal epithelium.

### Concordant increase of goblet cells and suppressive macrophages distinguish the COVID-infected nasal epithelium

We then assessed the differentially expressed genes between COVID-positive and COVID-negative samples within the epithelial **(Fig. 2A, left)** and immune **(Fig. 2A, right)** compartments. An expected increase in inflammatory and interferon-induced genes in both epithelial *(NFKBIA, C4B, IFITM2, ISG15, IFI6)* and immune compartments *(NFKBIA, C4B, IFITM1)* was observed in COVID-positive samples. However, we also found a co-enrichment of epithelial remodeling genes *(EFNA1, KRT6A/B/C, KRT7)* and immune suppressive macrophage genes *(CD163, C1QA, C1QB)*, suggesting a contrast in cell compositions. We next deconvolved cell populations **(Fig. 2B)** and found an enrichment of ionocytes and goblet/secretory cells in the epithelial compartment **(Fig. 2C, top)**, as well as higher macrophages and dendritic cells in the immune compartment **(Fig. 2C, bottom)**, in COVID-positive samples. Due to the minimal abundance of ionocytes **(Fig. 2B, top)**, we focused on the dynamics between macrophages and goblet cells.

**Figure 2.**
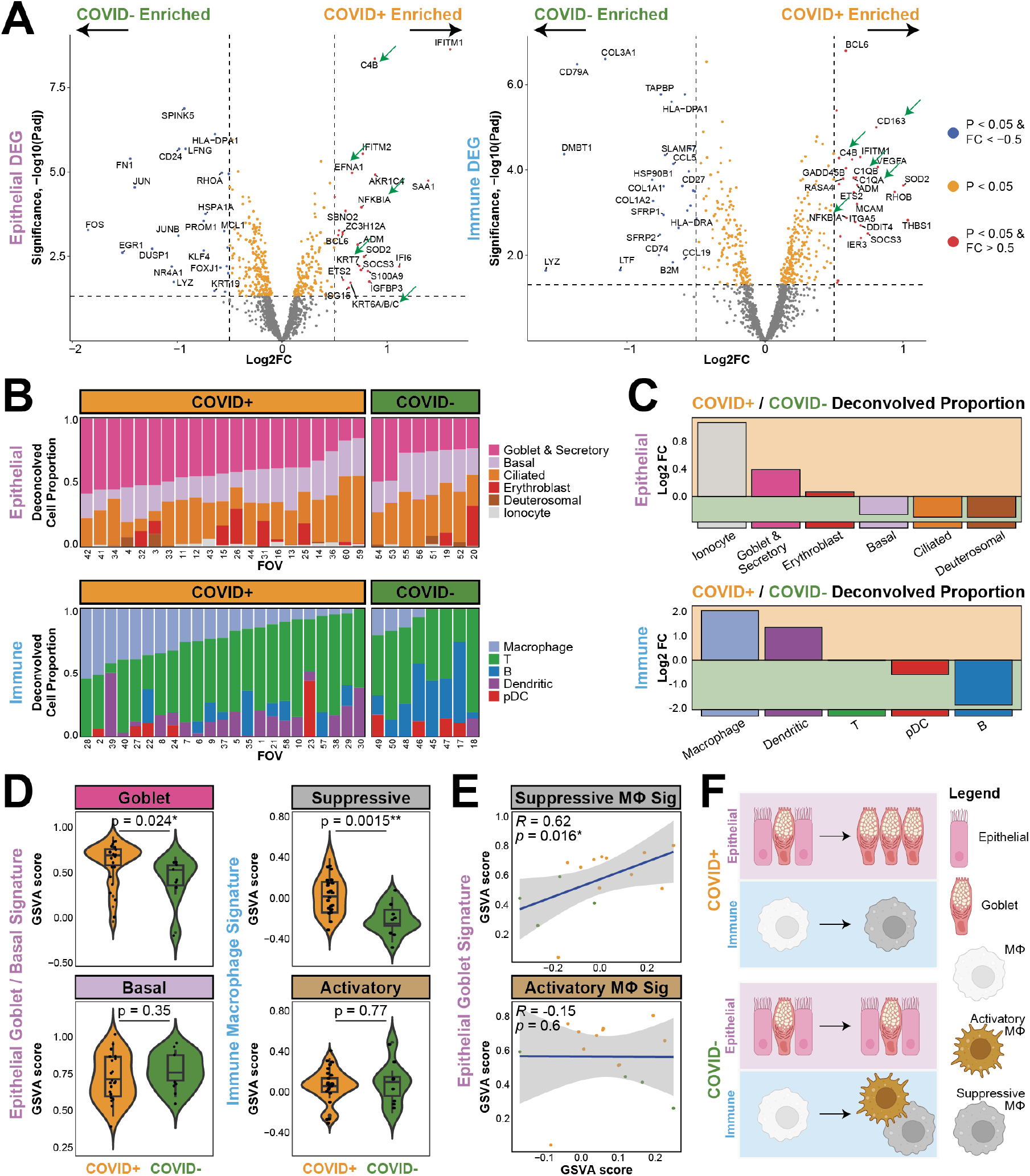
Distinct epithelial and immune cell compositions between COVID-positive and COVID-negative nasal tissues. **(A)** Volcano plots showing differentially expressed genes (DEG) between COVID-positive and COVID-negative tissues in the epithelial (left) and immune (right) compartments. Notice the green arrows that point to inflammatory, epithelial remodeling, and suppressive macrophage genes. **(B)** Relative proportions of deconvolved cell types **(Supp Table 1)** across COVID-positive and COVID-negative FOVs stratified by epithelial (top) or immune (bottom) compartments. **(C)** Log2 fold enrichment plot of deconvolved cell types between COVID-positive and COVID-negative FOVs stratified by epithelial (top) or immune (bottom) compartments. **(D)** Comparison of epithelial goblet/basal (left) and immune suppressive/activatory macrophage gene signatures **(Supp Table 1)** between COVID-positive and COVID-negative FOVs. A two-sided Wilcoxon rank sum test was conducted for each comparison (* *p ≤* 0.05, ** *p ≤* 0.01). **(E)** Spearman correlation of epithelial goblet gene signature with immune suppressive (top) or activatory (bottom) macrophage signature (* *p ≤* 0.05). Each dot represents one individual, and grey boundaries indicate 95% confidence interval. **(F)** Cartoon depicting the main differences in cell populations identified in COVID-positive and COVID-negative nasal epithelium.

These observations suggest that the COVID-positive nasal epithelium promotes a shift from progenitor (basal) to differentiated (goblet) states alongside macrophage polarization towards an immunosuppressive state, motivating a comparison of epithelial basal *vs*. goblet signatures as well as immune activatory *vs*. suppressive macrophage signatures. We found a significant increase in goblet and suppressive macrophage signatures in COVID-positive tissues, with no differences in basal cell or activatory macrophage signatures **(Fig. 2D)**. Furthermore, goblet cell signatures positively correlated with immunosuppressive, but not activatory macrophage signatures **(Fig. 2E)**. These findings support a functional relationship between the epithelial and immune compartments, through the concordant upregulation of goblet cells and suppressive macrophages, in the COVID-infected nasal epithelium **(Fig. 2F)**.

### Immune activation and IL13 expression links goblet cell and suppressive macrophages upregulation in the COVID-positive nasal epithelium

To determine the pathways that coordinate epithelial and immune remodeling, we identified the genes that were positively correlated with goblet and suppressive macrophage signatures in the COVID-positive nasal epithelium **(Supp Table 2)** and scored them against the Hallmark gene set collection (23). Inflammatory signatures were consistently the top features in both the epithelial and immune compartment, as well as increased metabolism (e.g. hypoxia and glycolysis), consistent with cellular energetic demands during viral infections (24) **(Fig. 3A)**. While inflammation was consistently elevated in COVID-positive tissues, further dissection of inflammatory-specific signatures revealed a dominant increase of acute inflammation and not type-2 responses specifically in the immune compartment **(Fig. 3B)**.

**Figure 3.**
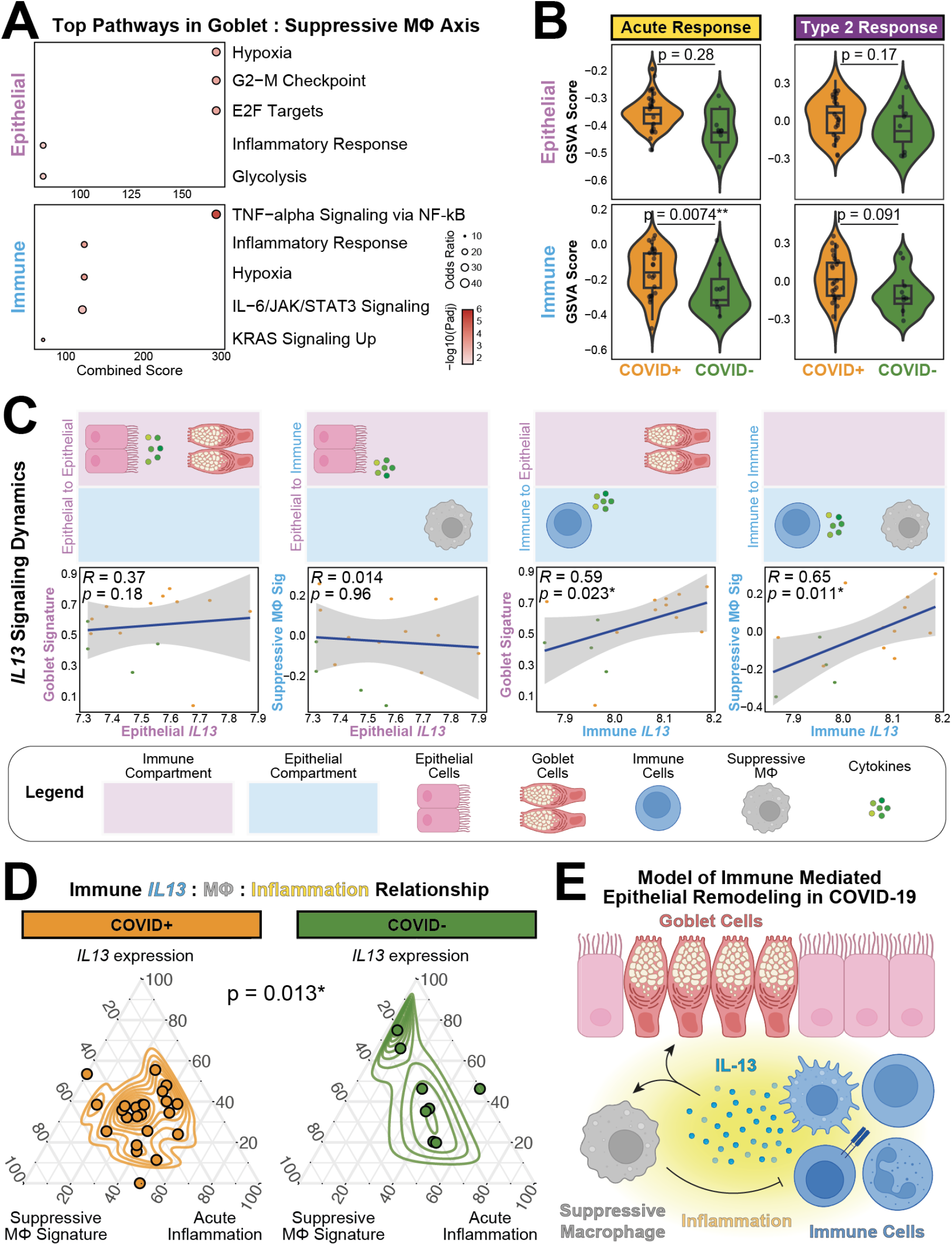
Expression of *IL13* in the immune compartment correlates with changes in immune and epithelial compositions. **(A)** EnrichR (21, 22) results for the top 5 Hallmark pathways (23) in COVID-positive epithelial (top) and immune (bottom) compartments using the genes positively correlated with both goblet and suppressive macrophage signatures. Scores for each pathway are in **Supp Table 2). (B)** Comparison of acute (left) and type-2 (right) inflammatory responses within epithelial (top) and immune (bottom) compartments between COVID-positive and COVID-negative tissues. A two-sided Wilcoxon rank sum test was conducted for each comparison (* *p ≤* 0.05, ** *p ≤* 0.01). **(C)** Spearman correlation of *IL13* expression levels with epithelial goblet or immune suppressive macrophage signatures (* *p ≤* 0.05). Each dot represents one individual, and grey boundaries indicate 95% confidence interval. Diagrams above each correlation indicate the regions where *IL13* and cell type signatures were analyzed. **(D)** Ternary plot illustrating the relationship between *IL13* expression, suppressive macrophage, and acute inflammation signatures in the COVID-positive and COVID-negative tissues. Each dot represents 1 FOV in the immune compartment, and contour lines indicate density distributions. A MANOVA test was conducted between COVID-positive and COVID-negative tissues (* *p ≤* 0.05). **(E)** Cartoon depicting the model of immune-mediated epithelial remodeling in the COVID-positive nasal epithelium.

The link between goblet cell differentiation from basal cells and macrophage polarization toward immunosuppressive subsets has been previously shown to be coordinated by soluble factors, including IL13 and IL4 (25, 26). We therefore searched for genes with expression levels that correlated with both goblet and macrophage signatures in our dataset **(Supp Table 2)** and identified an association of *IL13*, but not *IL4*, with both signatures. Notably, a previous study has also found an association between IL13, but not IL4, with COVID-19 disease severity (27). The accumulation of these knowledge motivated us to test the directionality of epithelial and immune remodeling by cross-correlating the expression of *IL13* with goblet and suppressive macrophage signatures. We found that only *IL13* in the immune compartment was positively correlated with both epithelial goblet and immune suppressive macrophage signatures **(Fig. 3C)**, suggesting that immune compartment-derived *IL13* is a key feature promoting epithelial remodeling during SARS-CoV-2 infection of the nasal epithelium.

These findings support a role for an active immune response in the COVID-positive nasal epithelium that involves suppressive macrophages, inflammation, and *IL13* expression, tied to an increase in goblet cells. A multivariate analysis of variance (MANOVA) test indicated that the composite profiles of these three signatures differed significantly between COVID-positive and COVID-negative groups. Consistent with this, visualization on a ternary plot revealed that COVID-positive samples occupied the centric region, reflective of the high contribution from all three axes, in contrast to COVID-negative samples **(Fig. 3D)**. Our spatial transcriptomics data support a model in which immune activation elevates *IL13* expression that leads to suppressive macrophage enrichment and maintain goblet cell formation in COVID-positive nasal epithelium **(Fig. 3E)**. We thus hypothesized that IL13-expressing cells spatially localize near the epithelial barrier and macrophages to coordinate immune-epithelial remodeling as a host response to viral infection.

### Single-cell spatial proteomics to interrogate mechanisms of nasal immune-epithelial remodeling in the COVID-positive nasal epithelium

To test this hypothesis, we applied the CODEX singlecell spatial proteomics platform (19) on 12 nasal epithelial FFPE tissues derived from 6 COVID-positive patients, with a combined protein and nucleic acid staining approach (28, 29), to visualize a panel of 22 protein and RNA targets **(Fig. 4A & Supp Fig. 1)**. The panel was designed to include key phenotyping markers that identify epithelial and immune cell types, functional markers focused on immune dysregulation, as well as RNA probes targeting *IL13* to detect *IL13*-expressing immune cells due to the difficulty of staining soluble factors in FFPE tissues.

**Figure 4.**
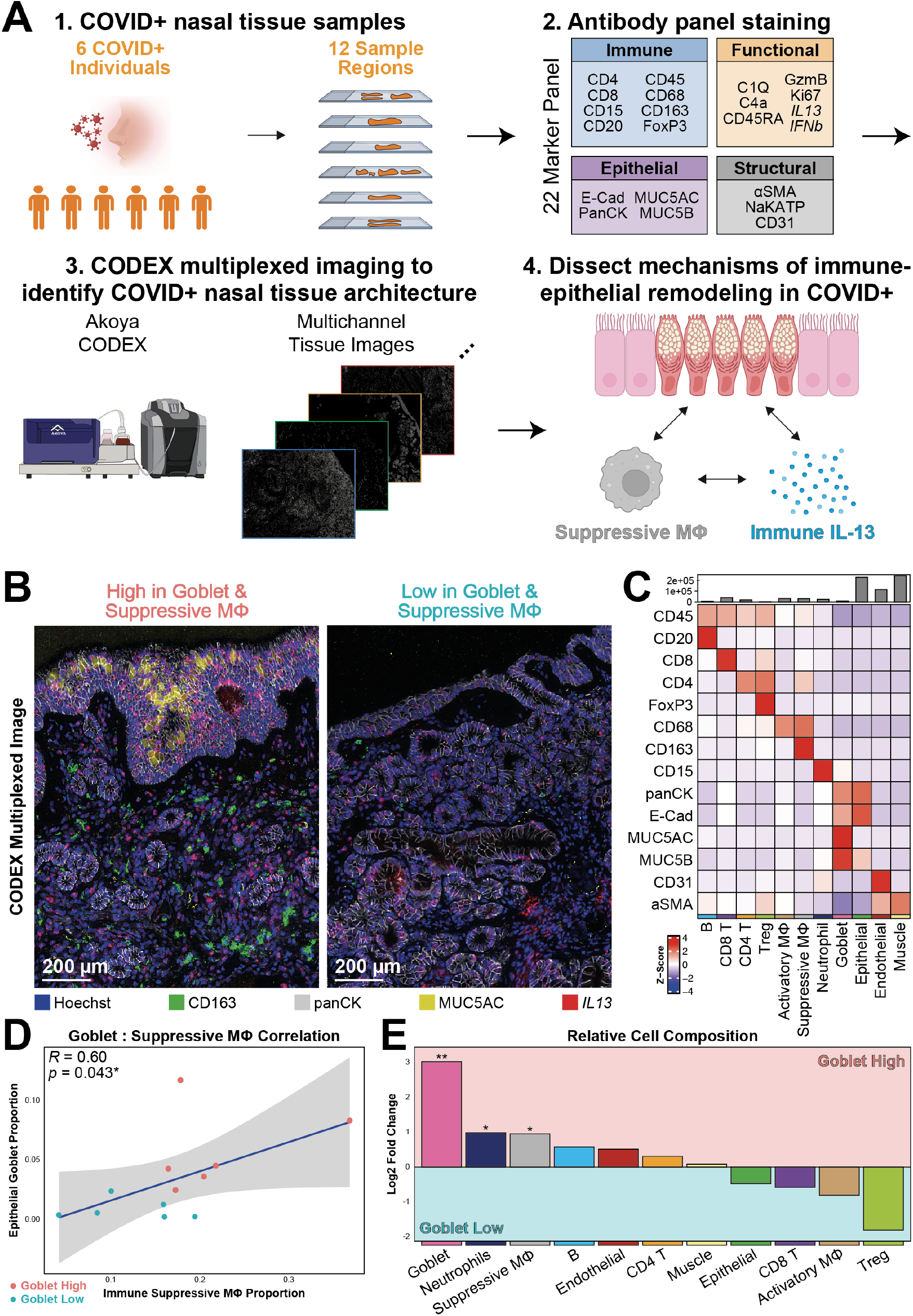
Application of CODEX single-cell spatial proteomics to dissect spatial mechanisms of immune-epithelial remodeling in the COVID-positive nasal epithelium. **(A)** Strategy to spatially dissect the COVID-positive nasal epithelium. (1) Tissues biopsies (n=12) from a cohort of COVID-positive patients (n=6) were (2) stained with a 22-plex antibody panel to enable (3) generation of multiplexed images that allowed for visualization of the tissue architecture at single-cell resolution, where upon (4) computational analysis revealed mechanisms promoting immune-epithelial remodeling in COVID-positive tissues. **(B)** Representative CODEX images of regions high in goblet cells and suppressive macrophages (left), in contrast to those low in goblet cells and suppressive macrophages (right), with markers for nuclei (Hoechst), suppressive macrophages (CD163), epithelial cells (panCK), goblet cells (MUC5AC) and *IL13* shown. **(C)** Relative z-score expression levels of phenotyping markers in this CODEX dataset. **(D)** Spearman correlation between epithelial goblet cell and immune suppressive macrophage proportions (* *p ≤* 0.05). Each dot represents one tissue, with the color indicating the Goblet High or Low group they are assigned to based on their goblet cell abundances. Grey boundaries indicate 95% confidence interval. **(E)** Log2 fold enrichment plot of annotated cell types between Goblet High and Goblet Low tissues. A two-sided Wilcoxon rank sum test was conducted for each comparison (* *p ≤* 0.05, ** *p ≤* 0.01).

Our cell type annotations **(Supp Fig. 2)** were stringently inspected by visually comparing to the original multiplexed images **(Fig. 4B)**, and by quantifying the expression of phenotyping markers to confirm their expected enrichment in their associated cell types **(Fig. 4C)**. During our visual inspection, we observed a contrast in the abundance of MUC5AC-positive goblet cells and CD163-positive suppressive macrophages, where some tissues were highly abundant in both while their presence was substantially lower in others **(Fig. 4B)**. Our results indicate a positive correlation between the proportion of suppressive macrophages and goblet cells **(Fig. 4D)**, orthogonally aligning with our spatial transcriptomics findings **(Fig. 2E)**. To test how this variability in goblet cell and suppressive macrophage abundance may be linked to immune-epithelial tissue microenvironments and remodeling, we next divided the tissues into two tissue groups (n=6 each) using a median cutoff based on their relative abundance of goblet cells and suppressive macrophages (**Supp Fig. 3A**). Further dissection of cell composition differences additionally revealed a significant i ncrease o f suppressive macrophages in the Goblet High group, as compared to inflammatory CD8 T cells and activatory macrophages in the Goblet Low group (**Fig. 4E**). These two groups were used henceforth to dissect spatial mechanisms underlying immune-epithelial dynamics.

### Tissue reorganization and spatial localization of IL13-positive CD4 T cells underlie nasal immune-epithelial remodeling

We first quantitatively assessed changes in overall tissue organization using two metrics that capture distinct properties of cellular composition. Shannon’s diversity index (30), which measures the diversity of cell types, was significantly h igher i n Goblet High compared t o Goblet Low tissues **(Fig. 5A)**. Ripley’s K function (31), which quantifies s patial c lustering, r evealed t hat c ells i n b oth tissue groups were aggregated relative to a random distribution, but Goblet High tissues were significantly less clustered than Goblet Low tissues **(Fig. 5B)**. Goblet High tissues thus exhibit a more heterogeneous and spatially dispersed cellular landscape, consistent with active immuneepithelial remodeling. We further detected lower inflammatory marker expression in Goblet High tissues **(Supp Fig. 3B)**, consistent with the compositional and spatial differences in tissue microenvironmental states observed above.

**Figure 5.**
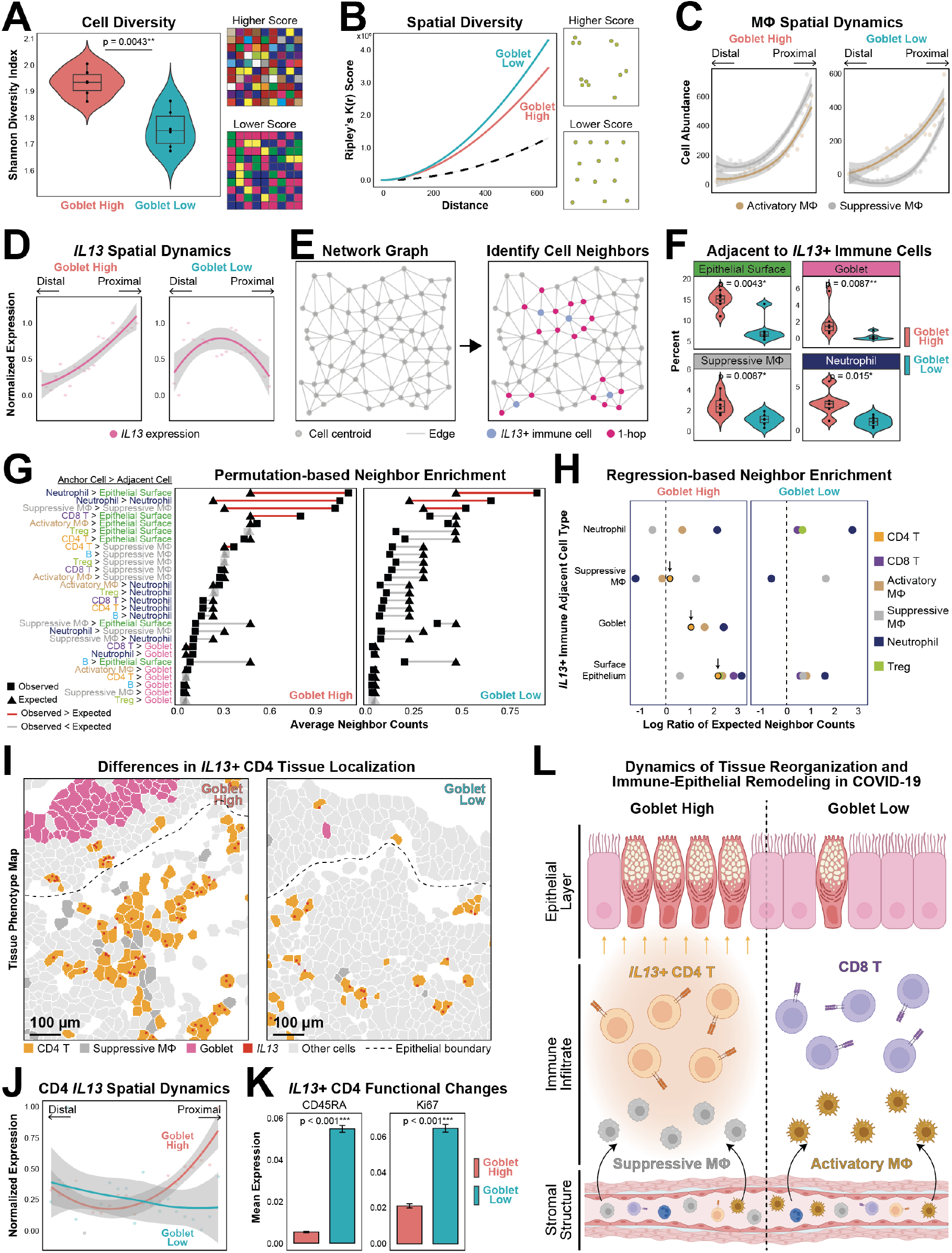
Spatial organization of *IL13*-positive CD4 T cells coordinates tissue remodeling in COVID-positive nasal epithelium. **(A)** Comparison of cell type diversity between Goblet High and Goblet Low tissues using the Shannon Diversity Index. A two-sided Wilcoxon rank sum test was conducted (** *p ≤* 0.01). **(B)** Comparison of cell spatial diversity between Goblet High and Goblet Low tissues using Ripley’s K function. The black line represents the expected diversity. **(C)** Differences in the infiltration of activatory and suppressive macrophages from surface-distal to surface-proximal layers in Goblet High (left) and Goblet Low (right) tissues. Each dot represents average abundance across tissues within a given spatial bin. **(D)** Differences in immune *IL13* expression from surface-distal to surface-proximal layers in Goblet High (left) and Goblet Low (right) tissues. Each dot represents average *IL13* expression across immune cells within a given spatial bin. **(E)** Cartoon of cell network graph visualization of the tissue to guide identification of *IL13*-positive immune cells and their adjacent cells. **(F)** Comparison of the abundance of cell types adjacent to *IL13*-positive immune cells between Goblet High and Goblet Low groups. A two-sided Wilcoxon rank sum test was conducted for each comparison (* *p ≤* 0.05, ** *p ≤* 0.01). Only the significant cell types are shown here, with the rest in **Supp Fig. 4B. (G)** Comparison of observed *vs*. expected counts of adjacent cells (the four significant cell types in **(C)**) around each anchor cell type (*IL13*-positive immune cells). The expected counts was generated over 1,000 iterations of randomized background permutations. Non-significant pairs are colored in grey. **(H)** Negative binomial regression results, with the predictor as anchor *IL13*-positive immune cells and the response as expected counts of adjacent cells (as in **(C)** and **(D)**, in Goblet High (left) and Goblet Low (right) tissues. Only statistically significant results (* *p ≤* 0.05) are shown. **(I)** Representative phenotype maps showing localization of *IL13* in CD4 T cells, goblet cells, and suppressive macrophages in Goblet High (left) and Goblet Low (right) tissues. To improve visual clarity, all other cells were greyed out with an epithelial boundary line manually drawn. **(J)** Relative expression of *IL13* in CD4 T cells from surface-distal to surface-proximal layers between Goblet High and Goblet Low tissues, with the other immune cells in **Supp Fig. 4C**. Each dot represents average *IL13* expression in CD4 T cells within a given spatial bin, and grey boundaries indicate 95% confidence interval. **(K)** Comparison of functional marker expression in *IL13*-positive CD4 T cells between Goblet High and Goblet Low tissues. Mean ± 1 standard error shown, with a two-sided Wilcoxon rank sum test conducted for each comparison (*** *p ≤* 0.001). **(L)** Cartoon depicting the proposed immune-epithelial remodeling dynamics in the COVID-positive nasal epithelium.

We next examined changes in the nasal tissue architecture relative to the surface epithelium by stratifying tissues into distance-based cellular neighborhoods, where regions were binned based on increasing spatial distance from this boundary **(Supp Fig. 4)**. Given the highly structured nature of the nasal epithelium that includes distinct epithelial immune, vascular, and stromal boundaries, we defined these cellular compartments by summarizing the composition of each cell’s nearest neighbors and clustering them based on similar compositional profiles **(Supp Fig. 5)**. Moving towards the surface epithelium, we observed a graded decrease in stromal cells accompanied by an increase in immune cells, with minimal changes in other compartments **(Supp Fig. 6A)**. This spatial pattern is consistent with the expected recruitment of immune cells towards the virus-infected surface epithelium.

There were key differences in immune spatial dynamics between the Goblet High and Goblet Low groups. While the relative compositions of most immune cells remained consistent **(Supp Fig. 6B)**, macrophages displayed distinct spatial dynamics, with preferential enrichment of suppressive macrophages in Goblet High and activatory macrophages in Goblet Low tissues **(Fig. 5C)**. These differences were consistent with spatial changes in immune *IL13* expression. In Goblet High tissues, it consistently increased toward the epithelial surface, but is spatially disorganized in Goblet Low tissues **(Fig. 5D)**. Despite these differences, the relative proportions of *IL13*expressing immune cells were similar between both tissue groups **(Supp Fig. 6C)**. This suggests that the spatial localization, rather than abundance, of *IL13*-expressing immune cells is associated with immune–epithelial remodeling dynamics.

To further investigate the spatial localization of *IL13* expressing immune cells, we next constructed a cell network graph to identify their immediate 1-hop neighbors **(Fig. 5E)**. We found that suppressive macrophages, goblet cells, neutrophils, and surface epithelial cells were more frequently adjacent to *IL13*-expressing immune cells in Goblet High tissues **(Fig. 5F)**, with no significant differences observed for the other cell types **(Supp Fig. 6D)**. These spatial associations support our model **(Fig. 3E)** that *IL13*-expressing immune cells are positioned to engage locally with macrophages and surface epithelium to drive immune-epithelial remodeling.

To evaluate if *IL13*-expressing immune cells were preferentially localized next to these cell types in Goblet High microenvironments, we applied a permutation test comparing the number of observed adjacent cells against an expected randomized background model, which was generated by shuffling t heir s patial p ositions w hile preserving the native locations of other cells (28). Although both Goblet High and Low tissues exhibited frequent *IL13*-centered interactions between neutrophil-surface epithelium, neutrophil-neutrophil, and suppressive macrophage-suppressive macrophage pairs, all other pairwise interactions were higher in Goblet High tissues with observed interaction frequencies closely matching the expected values, whereas in Goblet Low tissues, these interactions were consistently lower than expected **(Fig. 5G)**. These results demonstrate a spatial association between *IL13*-expressing immune cells and their neighboring cell populations in Goblet High tissues.

We next performed a regression analysis to assess which *IL13*-positive immune subtype was preferentially adjacent to these neighbors. Consistent with our permutation test, we found a greater diversity of *IL13*-expressing immune cells adjacent to suppressive macrophages, goblet cells, and surface epithelial cells in Goblet High tissues. *IL13*-expressing CD4 T cells were notably present in higher frequencies adjacent to suppressive macrophages, goblet cells, and surface epithelium specifically in Goblet High tissues, but not in Goblet Low tissues **(Fig. 5H)**. These findings w ere v isually c orroborated w ith o ur C ODEX images, where *IL13*-positive CD4 T cells were often in close proximity to the surface epithelium, goblet cells, and suppressive macrophages in Goblet High, but much less in Goblet Low microenvironments **(Fig. 5I)**, supporting *IL13*-expressing CD4 T cells as a major cell population involved in coordinating *IL13*-mediated immune-epithelial remodeling.

Further dissection of CD4 T cell functional state revealed an increase in *IL13* expression toward the surface epithelium in Goblet High tissues, whereas those in the Goblet Low group remained relatively consistent across tissue depths **(Fig. 5J)**. This *IL13* spatial gradient is not observed across other immune populations **(Supp Fig. 6E)**, suggesting that CD4 T cells also progressively upregulate *IL13* as they localize to this tissue niche. In addition, *IL13*-positive CD4 T cells in Goblet High tissues displayed reduced expression of naïve (CD45RA) and proliferative (Ki67) markers compared to those in Goblet Low tissues, consistent with a more differentiated phenotype **(Fig. 5K)**. These findings support CD4 T cells as a major *IL13*-producing cell type that represent a distinct, spatially organized population that coordinate tissue remodeling.

Our findings thus far suggest a protective role for IL13 signaling in viral infection, leading to the hypothesis that the nasal epithelium of patients with more severe infection would exhibit increased IL13-responsive activity, goblet cell metaplasia, and mucus production. However, we were unable to test this hypothesis with our samples due to the lack of viral titer measurements or patient disease severity. We therefore leveraged an independent RNA sequencing dataset (15) of human nasopharyngeal swabs stratified by COVID-19 disease severity (15 COVID-negative, 14 mild/moderate COVID-positive, 21 severe COVID-positive). We found that the surface epithelium exhibited robust *MUC5AC* and modest *MUC5B* increment across disease stages **(Supp Fig. 7A)**, substantial upregulation of IL13-inducible and goblet cell metaplasia gene signatures **(Supp Fig. 7B)**, as well as elevated PAMP, damage, and stress response genes **(Supp Fig. 7C)**. This supports the spatial relationship of immune IL13 **(Figs. 5F-K)** alongside the notion that IL13-driven epithelial remodeling scales with disease severity, reflecting an adaptive host antiviral response to enhance antiviral barrier function.

Our spatial multi-omics analyses highlight a link between spatial reorganization of the immune microenvironment and epithelial remodeling in COVID-positive nasal tissues. Regions enriched in goblet cells displayed co-localization of *IL13*-expressing immune cells, particularly CD4 T cells, near the epithelial surface and adjacent macrophages. In contrast, regions with low goblet cell abundance lacked spatial organization of *IL13* expression, exhibited decreased localization of *IL13*-positive immune cells to the epithelial interface, and were instead abundant in inflammatory immune infiltrates **(Fig. 5L)**. Given how this variation was observed within COVID-positive samples, the immune-epithelial remodeling may be driven by a combination of viral infection and local tissue contexts, such as the positioning and functional differences of CD4 T cells. We thus sought a mechanistic dissection into how IL13 directly reshapes epithelial barrier properties and modulate antiviral responses.

### IL13 induces compositional, morphological, and secretory remodeling of the nasal epithelium

To directly assess how IL13 drives epithelial remodeling, we utilized a primary human nasal epithelial air–liquid interface (ALI) model (32) to examine the morphological and functional effects of recombinant IL13 stimulation on the epithelial barrier. While the effects of IL13 on the airway mucosa, particularly its impact on MUC5AC-positive cells, are well recognized (33–35), the cellular-level dynamic changes of secretory cells during IL13-induced mucosal remodeling remain incompletely understood. Here, we treated ALI-cultured human nasal epithelial cells (HNEs) with purified IL13 (10ng/mL) added to the basal medium to mimic the effects of IL13-secreting CD4 T cells on neighboring epithelial cells, and fixed t he cultures at 0, 3, and 6 days after treatment. We then performed immunofluorescence staining to visualize the MUC5AC- and MUC5B-positive goblet and secretory cells respectively, the two major mucus-producing cell populations in the nasal epithelium. In addition, acetylated tubulin (ACTUB) was used as a marker to identify ciliated cells, and phalloidin staining was used to delineate cell boundaries **(Figs. 6A-C)**.

**Figure 6.**
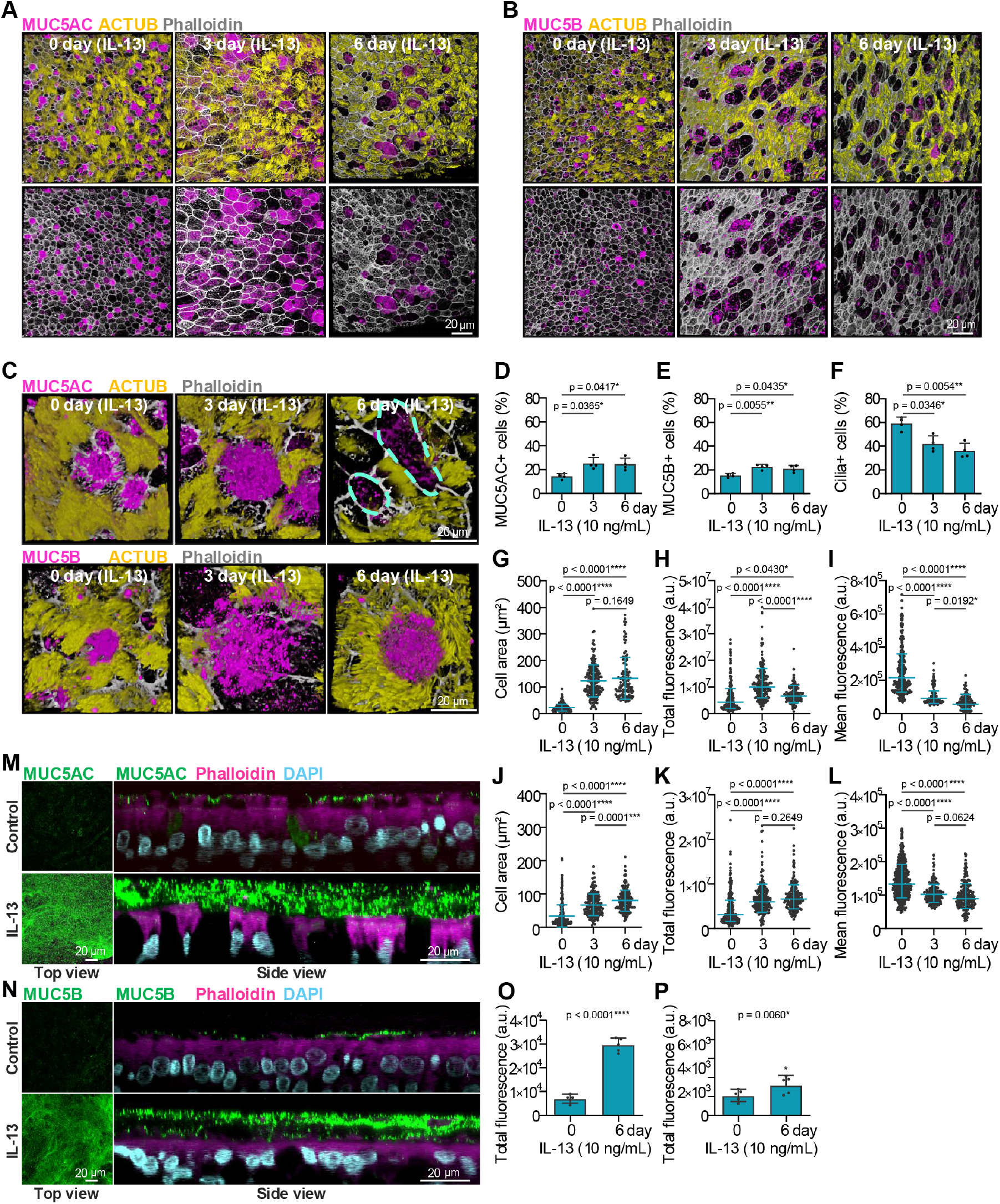
IL13 significantly remodels primary human nasal epithelium cultures. **(A-L)** IL13 remodeled the cellular composition and morphology of human nasal epithelium. HNE cultures were treated with IL13 in the basal medium for the indicated times, followed by fixation and immunofluorescence staining. Representative IF images are shown for MUC5AC (A & C), MUC5B (B & C), the cilia marker acetylated α-tubulin (ACTUB), and phalloidin to delineate cell boundaries. **(D-F)** Quantification of the percentages of MUC5AC-positive, MUC5B-positive, and ciliated epithelial cells. Each dot represents one donor. **(G & J)** Cell area of MUC5AC-positive or MUC5B-positive cells, measured based on phalloidin-defined cell boundaries. Data were collected from four independent donors. **(H, I, K, L)** Quantification of total cellular fluorescence intensity of MUC5AC (H) and MUC5B (K), and mean cellular fluorescence intensity normalized to cell area for MUC5AC (I) and MUC5B (L). Each dot represents one donor. **(M-P)** IL13 altered the composition of the apical mucus layer. Representative IF images are shown for MUC5AC (M) and MUC5B (N) in apical mucus following IL13 treatment. Quantification of MUC5AC (O) and MUC5B (P) fluorescence intensity in apical mucus is shown. **(D-L, O, P)** Error bars indicate mean ± SD (* *p ≤* 0.05, ** *p ≤* 0.01, *** *p ≤* 0.001, **** *p ≤* 0.001). (D–L) Paired one-way ANOVA with Tukey’s multiple-comparisons test. (O & P) Paired two-tailed Student’s t test. Representative fluorescence images were similarly observed in HNEs derived from four additional healthy donors.

**Figure 7.**
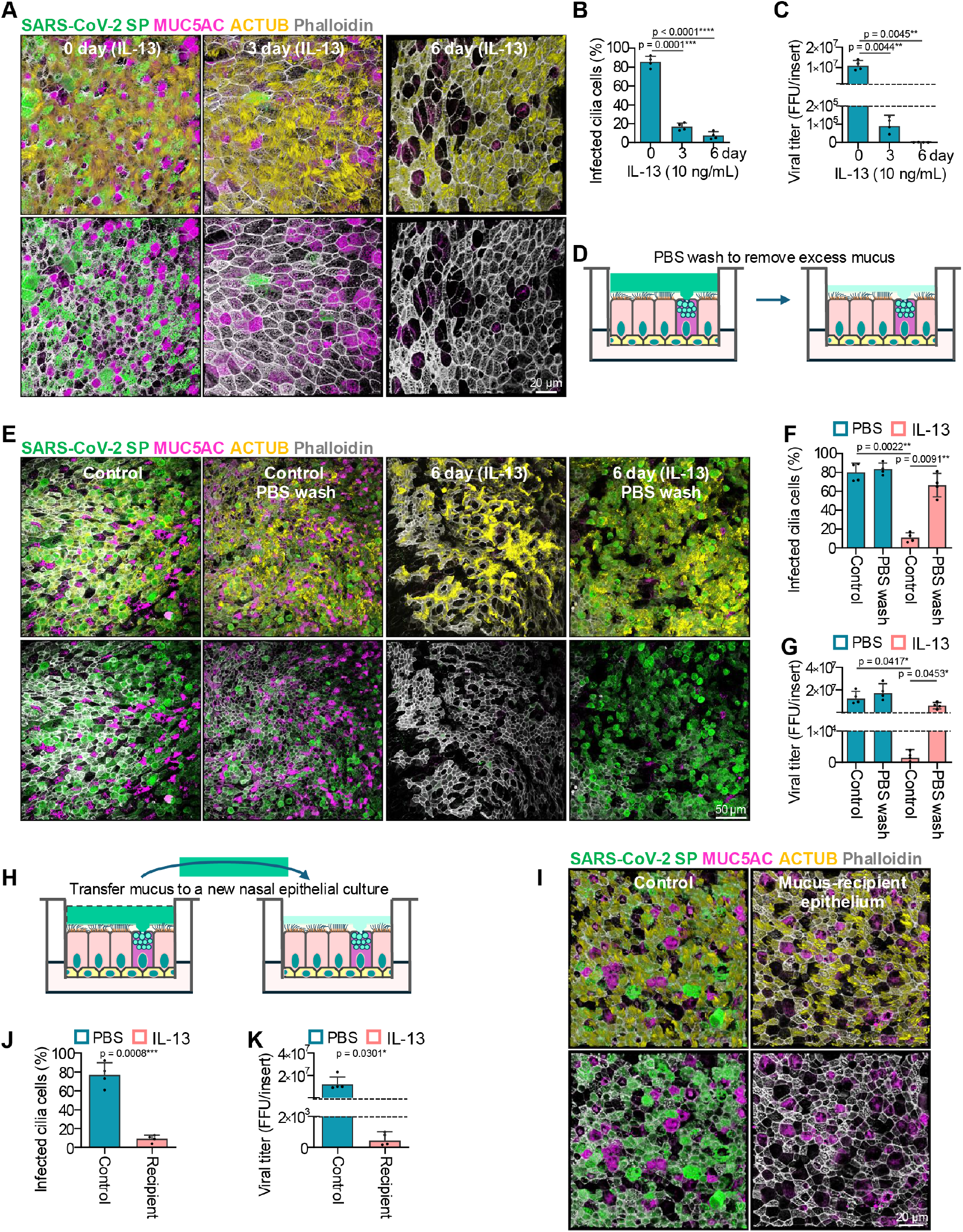
IL13-remodeled mucus effectively restricts viral entry. **(A-C)** IL13-remodeled nasal epithelium markedly suppressed SARS-CoV-2 infection. HNE cultures were treated with IL13 for the indicated times and infected apically with SARS-CoV-2 (MOI 0.1, 10 µL). At 48hpi, apical virus was collected in 100 µL PBS for FFU assay, and cultures were fixed for immunofluorescence analysis. (A) Representative IF images are shown for SARS-CoV-2 spike protein (SP), MUC5AC, ACTUB, and phalloidin. (B & C) Quantification of the percentage of SARS-CoV-2-positive ciliated cells (B) and viral titer in apical mucus measured by FFU assay (C). **(D-G)** Removal of IL13-induced thick apical mucus restored SARS-CoV-2 infection. (D) After 6 days of IL13 treatment, apical mucus was removed by incubating the apical surface with 100 µL PBS for 10 min and then aspirating the liquid; this procedure was repeated three times, followed by SARS-CoV-2 infection (MOI 0.1) and downstream analysis. (E) Representative IF images are shown for SARS-CoV-2 spike protein, MUC5AC, ACTUB, and phalloidin. (F & G) Quantification of the percentage of SARS-CoV-2-positive ciliated cells (F) and viral titer in apical mucus by FFU assay (G). **(H-K)** Transfer of IL13-induced thick mucus conferred protection to untreated HNE cultures. (H) After 6 days of IL13 treatment, 10 µL PBS was added to partially liquefy the apical mucus, which was then transferred onto untreated HNE cultures over 2 consecutive days (10 µL per day). Recipient cultures were subsequently infected with SARS-CoV-2 (MOI 0.1) and analyzed as above. (I) Representative IF images are shown for SARS-CoV-2 spike protein, MUC5AC, ACTUB, and phalloidin. (J & K) Quantification of the percentage of SARS-CoV-2-positive ciliated cells (J) and viral titer in apical mucus by FFU assay (K). **(B, C, F, G, J, K)** Error bars indicate mean ± SD (* *p ≤* 0.05, ** *p ≤* 0.01, *** *p ≤* 0.001, **** *p ≤* 0.001). (B, C, F, G) Paired one-way ANOVA with Tukey’s multiple-comparisons test. (J & K) Paired two-tailed Student’s t test. Representative fluorescence images were similarly observed in HNEs derived from four additional healthy donors.

From days 0 to 3 after IL13 treatment, there was a significant i ncrease i n t he p ercentages o f b oth MUC5AC-positive cells and MUC5B-positive cells accompanied by a reduction in the percentage of ciliated cells. These effects persisted at day 6 **(Figs. 6D-F)**, indicating that IL13 actively remodeled the nasal epithelium cellular composition during this period. Previous studies have shown that IL13 induces hypertrophy of MUC5AC-positive cells (33–35). Consistent with this, our data showed that IL13 significantly increased the size of MUC5AC-positive cells by day 3 **(Fig. 6G)**, accompanied by an increase in total intracellular MUC5AC fluorescence intensity **(Fig. 6H)**. By day 6, however, although MUC5AC-positive cells remained enlarged, their total intracellular MUC5AC fluorescence had decreased markedly **(Fig. 6C, top row & Fig. 6H)**, suggesting that substantial amounts of MUC5AC had been secreted from these cells in response to IL13. This interpretation was further supported by analysis of mean cellular fluorescence i ntensity, c alculated b y d ividing to-tal cellular MUC5AC fluorescence b y c ell a rea. Despite progressive enlargement of MUC5AC-positive cells, mean MUC5AC fluorescence intensity steadily declined from the onset of IL13 treatment **(Fig. 6I)**, indicating that MUC5AC secretion is a continuous process after initial IL13 stimulation.

Compared with our relatively established understanding of MUC5AC-positive cells, the effects of IL13 on MUC5B-positive cells remain less clear and, in some cases, controversial (36–40). Our data showed that under IL13 stimulation, MUC5B-positive cells also underwent significant hypertrophy at day 3 **(Fig. 6J)**, accompanied by a smaller but significant increase in total intracellular MUC5B fluorescence intensity **(Fig. 6K)** compared to MUC5AC. However, unlike MUC5AC-positive cells, MUC5B-positive cells remained enlarged at day 6 without a significant decline in total intracellular MUC5B fluorescence **(Fig. 6C, bottom row & Fig. 6K)**, suggesting that IL13 does not stimulate MUC5B secretion as strongly as MUC5AC. This is supported by the quantification of mean MUC5B fluorescence, which revealed a more modest decrease over time **(Fig. 6L)**. Consistent with these interpretations, our analysis of public sequencing dataset (15) revealed a higher magnitude of *MUC5AC* over *MUC5B* that scaled with disease severity and IL13 signaling **(Supp Fig. 7A)**. These findings help clarify the role of IL13 signaling in regulating MUC5B by showing that it contributes to MUC5B secretion, albeit at lower levels than MUC5AC. Consequently, IL13 stimulation shifts mucus composition toward a MUC5AC-skewed state.

Given that IL13 stimulation resulted in sustained MUC5AC and MUC5B secretion, we next asked whether this increased secretory activity translated into their accumulation within the overlying mucus layer, which would indicate a potential capability to alter epithelial barrier function. Immunofluorescence-based visualization **(Figs. 6M, 6N)** and quantification **(Figs. 6O, 6P)** of MUC5AC and MUC5B in the apical mucus layer showed that both mucins were significantly increased, with substantially pronounced increase in MUC5AC. Together, these findings demonstrate that IL13 begins to alter the morphology of mucus-secreting cells and the composition of the overlying mucus layer as early as day 3, and by day 6, it induces substantial remodeling of the nasal mucosa.

### IL13-induced mucus remodeling restricts viral entry in the nasal epithelium

Several studies have shown that IL13 treatment suppresses SARS-CoV-2 infection in airway epithelia, although the underlying mechanism remains incompletely understood (41–44). To determine whether the antiviral effect of IL13 is temporally linked to IL13-induced remodeling of mucus-secreting cells and the mucus layer, we pretreated ALI-cultured HNEs with IL13 for 0, 3, or 6 days before subjecting them to SARS-CoV-2 infection. To preserve the architecture of the apical mucus layer, SARS-CoV-2 was applied from the apical side in a small-volume inoculum (10µL, MOI 0.1). At 48 hours post-infection (hpi), progeny virus released into the apical mucus layer was collected for FFU assay, and the cultures were fixed for immunofluorescence analysis using antibodies against SARS-CoV-2 spike protein, ACTUB, and MUC5AC, together with phalloidin staining.

We found that IL13 pretreatment for either 3 or 6 days significantly suppressed SARS-CoV-2 infection in ciliated epithelial cells, with a stronger inhibitory effect observed after longer treatment **(Figs. 7A-C)**. Notably, this temporal pattern closely paralleled the kinetics of IL13-induced mucosal remodeling **(Fig. 6)**. Previous studies have suggested that IL13 may alter the expression of SARS-CoV-2 entry factors such as ACE2 and TMPRSS2 or modulate epithelial interferon responses (41–44). In our HNE model, qPCR analysis showed that IL13 caused no significant change in *ACE2* and *TMPRSS2* expression **(Supp Fig. 8A)**. In addition, IL13 treatment did not affect polyI:C-induced interferon production **(Supp Fig. 8B)**.

Given the pronounced effects of IL13 on the physical properties of the apical mucus layer, we hypothesized that the thickened mucus may act as a physical barrier that limits viral entry (38, 45, 46). To test this, we infected HNEs with SARS-CoV-2 at a high MOI (MOI 5) and fixed the cultures at 18hpi, thereby focusing on a single viral replication cycle and minimizing the potential contribution of IL13 to viral egress or secondary spread. Under these conditions, most ciliated cells in control cultures were infected by SARS-CoV-2, whereas the number of infected ciliated epithelial cells was markedly reduced in IL13-treated HNEs **(Supp Fig. 7C)**. These results lead to the hypothesis that an IL13-altered physical barrier impedes viral entry.

To directly test this hypothesis, we physically ablated the apical mucus layer from both control and IL13-treated HNEs by washing the apical surface three times with 100µL PBS for 10min each. The cultures were then infected with SARS-CoV-2 (MOI 0.1), and viral infection and apical viral output were analyzed at 48hpi **(Fig. 7D)**. Strikingly, removal of the apical protective layer from IL13-treated HNEs significantly increased both the number of infected ciliated epithelial cells and the amount of virus released into the apical mucus, restoring infection to levels close to those observed in control HNEs **(Figs. 7E-G)**. We next performed the reciprocal experiment by transferring the thick mucus layer from IL13-treated HNEs onto untreated control HNEs **(Fig. 7H)**. Remarkably, recipient cultures also acquired protection against SARS-CoV-2 entry, showing significantly reduced infection **(Figs. 7I-K)**. Together, these findings support a role for IL13 in protecting nasal epithelial cells from SARS-CoV-2 infection, in part through thickening of the apical mucus barrier to restrict viral entry.

## Discussion

Through *in situ* dissection of cross-sectional nasal tissues, our study uncloaks a spatially organized immune-epithelial microenvironment that promotes epithelial remodeling upon viral infection. In the context of SARS-COV-2 infected nasal tissues, IL13-producing immune cells, particularly CD4 T cells, localize to the surface epithelium and tissue macrophages to induce the formation of goblet cells and suppressive macrophages. This interaction is spatially restricted rather than uniform and diffuse across the tissue, a crucial insight given that regions lacking epithelial remodeling do not exhibit spatial patterns of IL13 expression despite showing a similar overall abundance of IL13-producing cells. Our findings thus highlight that spatial localization is also an important determinant of epithelial remodeling in the infected nasal epithelium beyond immune activation alone, particularly given the heterogeneity of goblet cell abundance within COVID-positive patients and other respiratory viral infections (47). This spatial heterogeneity may help explain why protective epithelial responses, such as mucus production, are not uniformly observed across infection (48).

Our findings extend current understanding of nasal tissue responses during COVID-19 infection. Previous studies, primarily based on nasopharyngeal swabs or airway washes, have independently reported transient inflammatory monocyte infiltration (15, 16), associations between IL13 signaling and disease severity (27, 49, 50), and presence of immunosuppressive M2-like macrophages (51– 53). By preserving the native tissue architecture, we show that these processes are not independent, but instead are spatially organized and converge at mucus production in goblet cells, a central component of upper airway antiviral defense (47). Our findings of the co-localization of *IL13*-producing CD4 T cells with epithelial and immune compartments provides a spatial framework that links immune signaling to epithelial remodeling. Building on these findings, functional experiments using a primary HNE ALI multi-cellular model demonstrate that IL13 alone is sufficient to drive significant morphological and functional epithelial remodeling capable of mounting antiviral defense *in vivo*. By integrating spatial analysis with functional validation, we directly link localized IL13 signaling at the epithelial surface to epithelial remodeling and mucus-mediated barrier protection, providing a mechanistic framework for how immune–epithelial interactions translate into antiviral function in human tissues.

Consequently, we were able to identify new insights **(Figs. 6–7)** beyond the well-established roles of IL13. By quantifying epithelial morphology across a temporal scale, we show that IL13-induced hypertrophy of MUC5AC-positive cells reflects a dynamic transition from intracellular mucus accumulation to sustained secretion, rather than a static endpoint. We clarified the effects of IL13 on MUC5B-positive cells by demonstrating similar hypertrophic and secretory responses, albeit to a lesser extent than MUC5AC. We also resolved an outstanding question regarding the mechanism by which IL13 suppresses SARS-CoV-2 infection by providing direct functional evidence that the remodeled mucus barrier with elevated MUC5AC and MUC5B, induced by IL13, itself mediates viral restriction, as its removal restored infection whereas its transfer conferred protection without modulating receptor expression or interferon-mediated responses as purported previously (41–44).

The relationship between IL13, goblet cells, and suppressive macrophages reflects a canonical type 2 immune response involving Th2 CD4 T-cells. In this study, we detected both type-1 and type-2 like signatures from our transcriptomics dataset **(Fig. 3B)** and observed the prevalence of *IL13*-expressing CD4 T cells **(Figs. 5H-K)**. In viral diseases such as COVID-19, inflammatory responses are expected to dominate (15, 16), but studies have also reported Th2 signatures within a Th1-dominant T-cell landscape (54). IL13 also does not always adhere to the Th2 vs. Th1 classification, as it has been detected in Th1 cells upon antigen exposure in mice (55) and implicated in immunoglobulin development post-vaccination (56). This coexistence warrants future investigation into the spectrum of T-cell functional states during viral infection of nasal airway.

A limitation of this study is the limited and difficult-to-procure sample size of nasal FFPE tissues and primary HNE cultures that are unable to support immune cells, and thus may not fully capture the range of patient-to-patient or donor-to-donor biological variability and the *ex-vivo* immune-epithelial interactions. Another limitation is the lack of viral titer and disease severity annotations in our samples. However, these limitations were mitigated by the integration of complementary spatial modalities, public sequencing datasets on independent cohorts, and functional validation in a physiologically relevant primary HNE model, supporting the generalizability of our findings. The multi-scaled analysis approach at progressively higher spatial resolution, spanning region-level transcriptomics to single-cell proteomics, enabled internal cross-validation across biological scales. Furthermore, to enhance the robustness and reliability of our HNE findings, each experiment was performed using nasal epithelial cells derived from at least 4 to 5 independent donors. This approach successfully circumvented an overreliance on sample size alone to establish consistency of findings. For instance, the correlation between immune-derived *IL13* and goblet cells observed in COVID-positive tissues in the GeoMx spatial transcriptomics dataset **(Fig. 3)** was corroborated with spatial localization and upregulation of *IL13* near goblet-enriched surface epithelia in the CODEX single-cell spatial proteomics dataset **(Fig. 5)** and upregulation of IL13 signaling and goblet metaplasia signatures from an independent patient cohort **(Supp Fig. 7)**. These compositional changes were then further recapitulated following direct *IL13* treatment in HNE culture **(Fig. 6)**. This exemplifies how our spatially resolved approach extends and refines prior studies by revealing a spatially organized immune–epithelial response that cannot be captured by non-spatial high-throughput modalities alone. We envision this framework to be broadly applicable across diverse disease contexts, illustrating how technological advances *in situ* and *in vivo* can be harmonized to drive deeper mechanistic insights.

In conclusion, we present a detailed dissection of the underlying immune and epithelial dynamics driving tissue remodeling in the virus-infected nasal epithelium. Our approach extends beyond most studies that rely on surface-level sampling such as swabs or washes, which is the only practical option for living patients, but fails to capture the deeper immune landscape that is only accessible through visualizing whole tissue cross-sections; this endeavor is only made possible through the exceptional initial efforts of our clinical partners and generosity of patient families in tissue donation. These findings u nderscore t he need to reframe how immune–epithelial interactions are studied and refine our understanding of the underlying biological mechanisms of *in situ* antiviral responses.

## Materials & Methods

### Tissue acquisition

SARS-CoV-2 infected sinonasal tissues were obtained during autopsy at the University of California San Diego and the University Hospital Basel. Non-SARS-CoV-2-infected sinonasal tissues from individuals with no clinical history, endoscopic, or radiographic signs of sinus disease were collected at the Stanford University Sinus Center. Tissues were processed in a similar manner as previously described (9, 10). All samples were acquired in accordance with the regulations of the Research Compliance Office u nder t he I nstitutional Review Board (University of California San Diego protocol ID #200485X, Stanford University protocol ID #18981) and approval by the ethics commission of Northern Switzerland (EKNZ study ID #2020-00969). All patients, or their relatives, consented to the use of tissue for research purposes.

### GeoMx data acquisition

Tissues were prepared according to the official N anostring G eoMx-NGS R NA Manual Slide Preparation protocol (Nanostring MAN-10150-01) as perfomed perviously (3). In brief, tissues were deparaffinized f or 3 0min a t 6 0°C b efore w ashing t hrice in xylenes at 5min each, twice in 100% EtOH at 5min each, once in 95% EtOH at 5min, and once in 1X PBS at 1min. Heat-induced epitope retrieval was then applied at 99°C for 10min in Tris-EDTA retrieval buffer (eBioscience 00-4956-58). After tissues were cooled, proteinase K (0.1g/ml) was next applied for 5min at 37°C, rinsed with 1X PBS, and fixed with 10% neutral buffered formalin (EMS Diasum 15740-04) for 5min at room temperature; fixation was later quenched using a 0.1M Tris and 0.1 M Glycine buffer for 5min and followed by a 5min wash in 1X PBS. The Human CTA detection probe cocktail was then applied to the tissues and incubated overnight (18 hrs) at 37°C. After probe hybridization, tissues were washed twice in Stringent Wash Buffer (2X SSC in 50% formamide) at 5min each. Buffer W (Nanostring 121300313) was then applied to the tissues for 30min followed by staining with the Solid Tumor TME Morphology Kit (GMX-RNA-MORPH-HST-12 containing anti-CD45, anti-panCK) for 1hr at room temperature. Tissues were then washed twice in 2X SSC at 5min each, stained with 500nM SYTO13 for 15min, and then loaded on the GeoMx. Capture ROIs were focused in areas with immune aggregation and epithelium on tissue apical surface. The CD45-positive immune and panCK-positive epithelial masks were finalized with agreement of at least two investigators.

### GeoMx data processing

After sample collection, the NanoString NGS library preparation kits were used to index each ROI with a unique Illumina i5 x i7 dual-indexing system according to the official Nanostring GeoMx-NGS RNA (Nanostring MAN-10153-01). Briefly, collected samples was mixed with PCR master mix and i5 and i7 primers for PCR amplification; the reaction conditions were 37°C for 30min, 50°C for 10min, 95°C for 3min, 18 cycles of 95°C for 15s each then 65°C for 60s then 68°C for 30s, and final extension of 68°C for 5min. Afterwards, the PCR product was purified with two rounds of AMPure XP beads (Beckman Coulter A63881) at 1.2X bead-to-sample ratio and paired-end sequenced on a NextSeq550. The GeoMx barcodes were then mapped and quantified using the default GeoMx analysis software, followed by batch effect correction and normalization as performed previously (57, 58). The normalized count table is in Zenodo.

### GeoMx data analysis

#### Principal component analysis

Dimensionality reduction was applied to all 1800 genes. The first two principal components were then used for visualization of sample clustering by tissue compartment and viral status.

#### Gene enrichment heatmap

Differential expression analysis was performed by comparing the magnitude and significance of each gene between the two conditions using Wilcoxon rank-sum tests (with Benjamini-Hochberg correction) and log_2_ fold change estimates. This analysis was run twice, first comparing immune to epithelial then comparing COVID-positive to COVID-negative, after which significant genes unique to each comparison were selected and visualized on a z-score heatmap.

#### Gene signature enrichment

Gene signatures were obtained from Molecular Signatures DataBase (MSigDB) (59) or prior resources (15, 60) and scored using Gene Set Variation Analysis (GSVA) (7). These genes were used to score goblet (*KRT7, CXCL17, F3, AQP5, CP, MUC5AC, KRT18, PTGS2*), basal (*KRT5, KRT15, TP63*), suppressive macrophages (*ARG1, IL10, CD163, PDCD1LG2, CD274, MARCO, CSF1R, IL1RN, IL1R2, IL4R, CCL4, CCL13, CCL20, CCL17, CCL18, CCL22, LYVE1, VEGFA, VEGFB, VEGFC, VEGFD, EGF, CTSD, TGFB1, TGFB2, TGFB3, MMP14, MMP19, MMP9, CLEC7A, WNT7B, TNFSF12, TNFSF8, CD276, MSR1, FN1, IRF4*), and activatory macrophages (*TNF, IL6, CD86, IL1B, MARCO, CD80, CXCL9, CXCL10, IL1A, CCL5, CD40, IDO1, CCR7*) respectively.

#### Differential gene expression

Limma (61) was implemented using the default parameters, and significance values were Benjamini-Hochberg corrected afterwards prior to visualization on a volcano plot.

#### Cell proportion deconvolution and composition

CIBER-SORT (62) was implemented using a reference resource (15) to deconvolve cell proportions, which were then used to calculate log_2_ fold change values accordingly.

#### Correlation analysis

The variables to correlate (e.g. GSVA scores or *IL13* expression across different cell populations) were performed using Spearman’s rank correlation.

#### Pathway enrichment identification

Genes f rom immune and epithelial regions were first c orrelated t o b oth goblet and macrophage signatures, and only those that were positively correlated and statistically significant t o both were used for EnrichR (21, 22) scoring using Hallmark pathways (23). The top 5 pathways were then filtered by adjusted p values and ranked by a combined score metric (ln(p) x z-score) for visualization. Correlated genes and pathway scores are in **Supp Table 2**.

#### Ternary plot

GSVA scores of suppressive macrophages and acute inflammation, a s w ell a s t he e xpression of *IL13* in each immune ROI, were first r escaled t o [0-1], from which a composite score was calculated and used to normalize such that the relative contributions of each summed to 1 within the immune ROI. A MANOVA test was also applied using the original non-normalized expression values, and significance was assessed using Pillai’s trace statistic with an alpha level of 0.05.

### CODEX antibody conjugation

Antibody conjugation was performed using a modified version o f (63). Briefly, carrier free antibody was concentrated using a 50kDa filter (MilliporeSigma UFC5050BK) and reduced using 0.9µM TCEP (MilliporeSigma C4706-10G) for 15-30min at room temperature. The reduction was quenched twice using a buffer containing 1mM Tris pH 7.5, 1 mM Tris pH 7.0, 150mM NaCl, 1mM EDTA, & 0.02% NaN_3_. Lyophilized maleimide oligos were resuspended in the same buffer supplemented with 250mM NaCl, and incubated with the antibody at a 2:1 mass ratio in 37°C for 1hr. The conjugated antibody was then washed thrice with a buffer containing 1X DPBS, 0.9M NaCl, & 0.02% NaN_3_ using a 50kDa filter and quantified using a NanoDrop (ThermoFisher ND-2000). Candor antibody stabilizer (ThermoFisher NC0414486) containing 0.2% NaN_3_ was then added to the antibody and stored at 4°C. Each conjugated antibody was carefully titrated before the actual study. Details regarding the antibody clones, vendors, barcodes, and titers are in **Supp Table 4**. More information on antibody selection and validation are in (64–66).

### CODEX data acquisition

Prior to tissue sectioning, glass coverslips (VWR 48366-067) were first silanized using Vectabond (Vector Laboratories SP-1800-7) according to manufacturer instructions. Briefly, coverslips were rinsed with ddH_2_O, soaked in acetone for 5min, transferred into 1:50 Vectabond diluted in acetone at 1:50 for 10min, and washed thrice in ddH_2_O. Any remaining liquid was gently dabbed clean using a KimWipe, air-dried at room temperature, and stored in a cool, dry place. After tissue sectioning, a PANINI-based implementation (28, 29) of RNAscope (67) was applied. In brief, tissues were de-paraffinized for 1hr at 70°C before washing twice in xylenes at 10min each, thrice in 100% EtOH at 3min each, twice in 95% EtOH at 3min, once in 80% EtOH at 3min, once in 70% EtOH at 3min, and thrice in ddH_2_O at 3min each. Heat-induced epitope retrieval was then applied at 97°C with a Target Retrieval Solution (Agilent S236784-2). After tissues were cooled, the coverslip edges were circled with a PAP pen (Vector Laboratories H-4000), blocked with 0.3% H_2_O_2_ at 40°C for 20min, followed by avidin block and biotin block at room temperature for 30min each; two ddH_2_O rinses were performed in between the blocks. C2 *IL13* probes (ACDbio 586241-C2) were then diluted 1:50 in C1 *IFN*β probes (ACDbio 417071) and hybridized overnight (18h) in 40°C. The following day, the RNAscope Multiplex Fluorescent Reagent Kit v2 (ACDBio 323100) was used for amplification as instructed, with the exception of using tyramide signal amplification reaction to deposit biotin (Akoya NEL749A001KT) and digoxigenin (Akoya NEL748001KT) haptens at C1 and C2 respectively. Tissues were then used for CODEX staining following previously described methods (63). In brief, tissues were blocked using a buffer containing 5% donkey serum (MilliporeSigma D9663-10ML), 0.05% NaN_3_, 50µg/mL mouse IgG (MilliporeSigma I5381-10mg), 50µg/mL rat IgG (MilliporeSigma I4141-10mg), 500µg/mL sheared salmon sperm DNA (ThermoFisher AM9680), and 50nM oligonucleotide block (Invitrogen AM9849) in 1X TBS-T while photobleaching for 2h prior to overnight incubation at 4°C with the antibody cocktail **(Supp Table 4)**. The next day, tissues were washed thrice in a buffer containing 2.5mM EDTA, 0.25% BSA, 0.02% NaN_3_, 250mM NaCl, 61mM

Na_2_HPO_4_, 39mM NaH_2_PO_4_ in 0.5X DPBS at 5min each, fixed with 1.6% PFA (EMS Diasum 15740-04) twice at 5min each, rinsed thrice in 1X PBS, fixed in ice-cold MeOH for 5min, rinsed thrice in 1X PBS, fixed i n 4 µ g/µL BS3 (ThermoFisher 21580) twice at 10min each in the dark, and finally rinsed thrice in 1 X PBS. Reporter plates were also prepared using a black 96-well plate (Corning 07-200-762), with containing a Cy3 and Cy5 fluorescent oligonucleotide and 1:600 of Hoechest (ThermoFisher H3570) corresponding to each imaging cycle; plates were sealed with alumnium film (ThermoFisher 1 4-222-342). The tissue coverslip was loaded to an inverted fluorescence microscope (Keyence BZ-X810) operated under a 20x/0.75 objective (Nikon CFI Plan Apo λ) while the reporter plate was loaded into a connected CODEX microfluidics. Each imaging cycle contained three channels, one for the nuclear stain and two markers on the Cy3 and Cy5 filter channels. For the first a nd l ast i maging c ycles, n o fluorescent oligonucleotides were included, enabling the Cy3 and Cy5 images in those cycles to serve as blank channels for background subtraction.

### CODEX data processing

The acquired CODEX images were stitched and background corrected using the Akoya Singer software. Cell segmentation was then performed using a local implementation of deepcell-tf version 0.12.2 (68, 69). The Hoechst channel was used for the nucleus, while the summation of NaKATP, CD31, CD45, a-SMA, pan-Cytokeratin, and CD15 was used for the membrane feature, with maxima_threshold = 0.4 and interior_threshold = 0.05 segmentation parameters for optimal capture of cell morphology. The single-cell feature expression matrix was then generated by adding the counts for each channel within individual cell segmentation mask boundaries and dividing by the single-cell mask area. The area-normalized values were then divided by the median Hoechst value for each tissue to ensure tissue-to-tissue consistency across different CODEX runs. The singlecell data was then arcsinh transformed using a cofactor of 150, and rescaled to a global [0-1]. The scaled single-cell expression of phenotyping markers were then iteratively clustered using a combination of PhenoGraph (70) and MAPS (71), followed by visualization on Mantis Viewer (72) to correct and confirm final cell annotations. The remaining unidentifiable cells that did not express any phenotyping markers were assigned as “Others”. As *IL13* signal were discrete puncta, Piscis (73) was applied to quantify the spots, which were then assigned to individual cells based on the cell segmentation mask. This normalized single-cell expression matrix is in Zenodo.

### CODEX data analysis

#### Identification of Goblet High and Low tissues

Two scoring approaches were applied: a rank-based score, defined as the average of the ranks for each proportion across tissues, and a proportion-based composite score, calculated by multiplying the proportion of goblet cells within epithelial cells by the proportion of suppressive macrophages within immune cells. Tissues were then stratified using a median cutoff based on the composite score, and both approaches resulted in the same stratifications. A final visual inspection was done to confirm the presence of contrasting goblet-high and goblet-low tissues.

#### Marker expression heatmap

Phenotyping marker expression levels were standardized (z-scored) across the annotated cell populations to assist confirmation of annotation specificity. Inflammatory marker expression were standardized across all tissues and cell populations, and the resulting z-scores were then averaged within each cell population to obtain a representative mean z-score per population to evaluate functional differences.

#### Correlation analysis

Goblet and suppressive macrophage cell proportions were Spearman correlated to validate the analysis in **Fig. 2E**.

#### Relative cell composition

Log_2_ fold change values were calculated using the global cell proportions between these two groups, with separate Wilcoxon rank-sum tests performed to assess statistical significance

#### Cell diversity

Shannon’s Diversity Index (30) (-*p*_*i*_ log *p*_*i*_) was used to determine cell diversity for Goblet High and Goblet Low tissues, after which they were compared using a Wilcoxon rank-sum test.

#### Spatial diversity

A convex hull was first generated around each tissue boundary to define the spatial window while also using cell centroid coordinates to construct a point pattern, from which Ripley’s K(r) Score and its confidence envelope were estimated using 99 Monte Carlo simulations and compared between tissue groups.

#### Distance neighborhood stratification

The surface epithelial layer was used as a reference boundary to partition tissues into spatial bins at 100 pixel intervals based on each cell’s minimum Euclidean distance to the reference **(Supp Fig. 3)**; for consistency across samples, downstream analyses were restricted to the lowest common maximum bin.

#### Spatial cell composition change

Local cell compartments were first defined by computing 20 nearest neighbors of each cell, and then k-means clustered with 5 centers **(Supp Fig. 4)**. The proportion of cells belonging to each cellular compartment, as well as the abundance of each immune cell type, was then summarized for each spatial bin identified from distance neighborhood stratification.

#### Spatial IL13 expression change

*IL13* spot counts were calculated in the immune compartment and normalized to the same [0-1] scale prior to visualization across spatial bins.

#### Sankey plot

Immune cells were grouped by cell type, *IL13* expression status (expressing *vs*. non-expressing), and tissue cluster (Goblet High or Goblet Low), from which cell counts were summarized for each combination of these categories and visualized accordingly.

#### Network graph projection of tissues

Tissues were represented as cell network graphs by applying Delaunay triangulation (74) on individual cell centroids, and two nodes are only connected if their Euclidean distance was less than or equal to 150 pixels.

#### Identification o f I L13-adjacent cells

*I L13*-positive immune cells were defined as anchor nodes, and all 1-hop neighbors were retrieved using the triangulated adjacency graph. These neighbors were then tabulated to quantify the percentage of neighboring cell compositions per tissue (excluding the anchor cells). Differences between tissue groups were assessed using Wilcoxon rank-sum tests.

#### Permutation-based neighbor enrichment

*IL13*-positive immune cell labels were randomly shuffled 1,000 times within each tissue to generate a null distribution of neighbor counts. Differences between observed and average expected counts were computed for each neighbor type, as well as the corresponding empirical p-values (proportion of permutations where expected counts are different observed counts) that were Benjamini-Hochberg corrected.

#### Regression-based neighbor enrichment

A negative binomial regression model was applied, where for each *IL13*-positive immune anchor, the counts of adjacent cells of interest were used as response variables, with the anchor cell type as the predictor. B cells were set as the baseline due to their stable but non-dominant abundance with consistent expression of *IL13*, with separate models fitted for Goblet High and Goblet Low tissues and significance values Benjamini-Hochberg corrected prior to evaluation.

#### Spatial dynamics of IL13 expression in immune cells

The average number of *IL13* spots in each immune population was quantified across each spatial bin within the separate tissue groups, and then normalized to the same [0-1] scale to aid visualization.

#### IL13-positive CD4 T cell marker analysis

The mean and standard error of CD45RA and Ki67 expression across CD4 *IL13*-positive cells were quantified and compared using a Wilcoxon rank-sum test.

### Analysis of public RNA sequencing dataset

The preprocessed .h5ad object from Ziegler et al. (15) was pulled from CELLxGENE (75), where patients were already grouped into disease severity categories based on the WHO disease scoring and was correlated with viral titer as showed by the authors. After filtering for epithelial cells, *MUC5AC* and *MUC5B* expression were averaged per patient within each severity group. Genes linked to IL13 downstream signaling activity (*CCL26, POSTN, CLCA1, SERPINB2, ALOX15, IL13RA2, ITLN1, DPP4, SOCS1, SOCS3, CCL17, CCL22, TGM2, CST1, SERPINB4*) and goblet cell metaplasia (*SPDEF, FOXA3, FOXM1, KLF4, TFF3, BPIFB1, AGR2, FCGBP, B3GNT6, ERN2*) were pseudobulked on a patient level and scored using GSVA (7). As these biological phenomena have overlapping genes, they were curated to be non-overlapping to avoid redundancy and improve interpretation of pathway activities. Differences between disease groups were then assessed using Wilcoxon rank-sum tests.

### Primary human nasal epithelial cell culture

Human nasal epithelial cells were obtained from healthy donors or patients undergoing nasal endoscopy or surgical polypectomy during diagnostic procedures for sinonasal or respiratory diseases at UT Southwestern Medical Center (UTSW). Nasal epithelial cultures were established using a well-established protocol routinely used in our laboratory (32). After informed consent was obtained (UTSW IRB# protocols STU-2023-1104 and STU-2019-1652), cells were collected by brushing the inferior turbinate of both nasal cavities. Samples were immediately transferred to the laboratory and dissociated for 10min at 37°C in a 1:1 mixture of proteolytic enzyme solution Accutase (ThermoFisher A1110501) and nonenzymatic cell dissociation solution (MilliporeSigma C5914). After gentle agitation, cells were pelleted, resuspended in basal cell proliferation medium PneumaCult-Ex Plus (STEMCELL 05040) supplemented with antibiotics, and seeded onto collagen-coated Transwell inserts (0.33cm^2^) at a density of 20,000 cells per insert, with medium added to both the apical and basal compartments. Once the cells reached confluence, the medium was replaced with PneumaCult-ALI medium (STEMCELL 05001) in the basal chamber, and the apical medium was removed to establish an air-liquid interface (ALI). Cultures were maintained at ALI for an additional >40 days to allow full differentiation of the nasal epithelium, including intact mucociliary transport, ciliated cells, basal cells, and multiple secretory cell types.

### Virus production and cell infection

SARS-CoV-2 BA.5 was a gift from Luis Martinez-Sobrido (Texas Biomedical Research Institute). Experiments involving SARS-CoV-2 were performed under Biosafety Level 3 (BSL-3) conditions, in accordance with protocols approved by the UT Southwestern Biosafety Committee. SARS-CoV-2 was propagated in Vero-hACE2-TMPRSS2 cells. RSV and GFP-RSV was amplified in HEp-2 cells, while PIV1, GFP-PIV1, and GFP-SeV were propagated in LLC-MK2 cells. Cells were grown in T75 flasks using an infection medium consisting of DMEM supplemented with BSA (Millipore-Sigma, A7979) and 7.5% sodium bicarbonate. The culture supernatants were harvested. The pooled viral stocks were passed through a 0.45µm filter (MilliporeSigma, SLHV004SL) to remove cellular debris. All viral stocks were further concentrated using Centricon Plus-70 PL-100 centrifugal filter units (MilliporeSigma, UFC710008) at 4,000rpm for 40–50min at 4°C to achieve working titers of approximately 106–107PFU/mL. Viral titers were determined by plaque assay on appropriate indicator cells, and viral identity was verified by Sanger sequencing of the viral genomes. To infect the cells, 10µL of virus suspension (MOI 0.1–5) was added to the top of the ALI-cultured HNEs.

### Viral titration and quantification

Viral titers were determined by plaque assay on Vero E6 cells. Confluent monolayers grown in six-well plates were incubated with serial dilutions of virus samples (250µL/well) at 37°C for 1h to allow adsorption. Following incubation, the inoculum was removed, and the cells were overlaid with 1% agarose (Invitrogen, A2576) prepared in MEM supplemented with 1% penicillin-streptomycin-glutamine (Gibco, 10378016). After incubation, cells were fixed w ith 4 % formaldehyde for 2h, the agarose overlay was removed, and plaques were visualized by staining with 0.1% crystal violet. Viral titers were expressed as plaque-forming units per milliliter (PFU/mL). Viral infectivity was further quantified by focus-forming assay (FFU) on Vero E6 cells in 96-well plates. Briefly, c onfluent mo nolayers we re in oculated wi th serial dilutions of virus for 1 h at 37°C . To ensure quantification within a single round of infection, the inoculum was replaced with fresh culture medium and cells were incubated 18-24 hours to allow for viral protein expression. Following incubation, cells were fixed with 4% formaldehyde, permeabilized with 0.1% Triton X-100, and blocked with 5% FBS in PBS. Infected cells were detected via immunofluorescence using virus-specific SARS-CoV-2 spike protein and nucleocapsid primary antibodies. After incubation with Alexa Fluor 488- or 568-conjugated secondary antibodies (Invitrogen), fluorescent f oci w ere v isualized a nd quantified using an EVOS FL Auto Imaging System (Invitrogen). Viral titers were expressed as FFU/mL.

### Antibodies for primary culture

Antibodies used include the following: mouse anti-SARS-CoV-2-nucleocapsid protein (clone E16C, ThermoFisher MA1-7403, 1:200), mouse anti-SARS-CoV-2-spike protein (clone 1A9, GeneTex GTX632604, 1:600), mouse anti-Acetylated alpha tubulin (clone 6-11B-1, Santa Cruz sc-23950, 1:600), rat anti-Alpha tubulin (clone YL1/2, ThermoFisher MA1-80017, 1:100), rabbit anti-MUC5B (polyclonal, Atlas HPA008246, 1:100), mouse anti-MUC5AC (clone 45M1, Abcam ab212636, 1:200), Alexa Fluor™ Plus 647 Phal-loidin (Invitrogen A30107), and Alexa Fluor™ Plus 405 Phalloidin (Invitrogen A30104).

### IF immunohistochemistry of cryosections

For cryosections, fully matured HNEs were fixed i n 4% PFA in 1X PBS for 30min, followed by three 5min washes with 1X PBS. HNEs were incubated with primary antibodies in the IF buffer (3% BSA and 0.4% saponin in PBS) overnight at 4°C, followed by 3 washes with the IF buffer. Samples were then incubated with fluorescent-labeled secondary antibody with IF buffer at room temperature for 4h, followed by a 5min incubation with DAPI in PBS at room temperature for 5min and 3 washes with IF buffer. Coverslips were mounted with Fluoromount-G (SouthernBiotech 0100-01) onto glass slides followed by image acquisition.

### Confocal microscopy

Fluorescence images were acquired using a Nikon AX R confocal microscope equipped with 405, 488, 561, and 640 nm lasers. High-speed live imaging was performed using an integrated ultra-high-speed camera capable of recording at up to 720 frames per second at 2048 × 16 pixel resolution. Samples were maintained under physiological conditions using the microscope’s incubation system. Post-acquisition image processing was performed using NIS-Elements (Nikon).

### Quantitative real-time PCR

RNA was extracted using the RNeasy Lipid Tissue Kit (QIAGEN) and cDNA was synthesized using M-MLV Reverse Transcriptase (Invitrogen, 28025-013). Quantitative real time PCR was performed using TaqMan Probes (Invitrogen) and the TaqMan Gene Expression Master Mix (Applied Biosystems, 4369016) in 96-well Micro Amp Optical reaction plates (Applied Biosystems, N8010560). Expression levels were normalized to the average expression of the housekeeping gene. Life Technologies TaqMan probes for human genes were ACE2 (Hs00222343_m1), TMPRSS2 (Hs00237175_m1), IFNB1 (Hs01077958_s1), IFNL1 (Hs00601677_g1), and GAPDH (Hs02786624_g1).

### Figure assembly

Figures were assembled using Adobe Illustrator **(Figs. 1-5 & Supp Figs. 1-7)** or Microsoft Powerpoint **(Figs. 6-7 & Supp Fig. 8)**, with necessary image objects for cartoon schematics exported from BioRender.

## Code and data availability

Code for analysis and data visualization are at https://github.com/SizunJiangLab/COVID-Nasal. GeoMx count table, CODEX images, and CODEX single-cell expression matrix are at https://doi.org/10.5281/zenodo. 20127355.

## ACKNOWLEDGEMENTS

We thank members of the Jiang and Wu labs for helpful discussions and Aniket Gad from Akoya Biosciences for CODEX technical support. S.J. is supported in part by NIH DP2AI171139, the Bill & Melinda Gates Foundation INV-002704, the Broad Next Generation Award, the Dye Family Foundation, and the Bridge Project, a partnership between the Koch Institute for Integrative Cancer Research at MIT and the Dana-Farber/Harvard Cancer Center. C.T.W. is supported by NIH R01AI184984 and the UT System Rising STARs award. Y.Y.Y. is a recipient of the Albert J Ryan Fellowship. A.T., M.S.M., G.P.N., D.R.M, and S.J acknowledge the integral early support from the Basel Research Center for Child Health. This work was supported in part by US Food and Drug Administration (FDA) Medical Countermeasures Initiative contracts 75F40120C00176 and HHSF223201610018C. This article reflects the views of the authors and should not be construed as representing the views or policies of the institutions that provided funding.

## AUTHOR CONTRIBUTIONS

S.J., I.T.L., T.N., Y.Y.Y., C.T.W., H.Y.C. conceived of the study. Y.Y.Y., H.Y.C., I.T.L., T.N., M.S.M., C.T.W., S.J. planned and performed the experiments. Y.Y.Y., H.Y.C., B.Z., Yang W., Yuchen W., H.Q., K.B.J. performed data analysis. S.C., C.H.Y., C.S.K., H.A.M., A.Z.M., W.W., J.V.N., G.P.N., D.R.M., A.T. contributed to reagents, clinical samples and/or technical expertise. Work by Yang W. was conducted during her tenure as a visiting member of the Jiang laboratory. Y.Y.Y., C.T.W., S.J. wrote the manuscript with input from all authors. S.J. & C.T.W. provided supervision and funding acquisition. All authors reviewed and approved the final manuscript.

