## Supplementary Figures for "Coordinated immune-epithelial dynamics in the nasal epithelium protect against respiratory virus infection"

[1]Center for Virology and Vaccine Research, Beth Israel Deaconess Medical Center, Harvard Medical School, Boston, MA, United States
[2]Program in Virology, Division of Medical Sciences, Harvard Medical School, Boston, MA, United States
[3]Department of Microbiology, UT Southwestern Medical Center, Dallas, TX, United States
[4]Department of Pathology, Stanford University, Stanford, CA, United States
[5]Department of Otolaryngology-Head and Neck Surgery, Stanford University School of Medicine, Stanford, CA, United States
[6]Department of Otolaryngology-Head and Neck Surgery, UT Southwestern Medical Center, Dallas TX, USA
[7]Department of Otolaryngology-Head and Neck Surgery, University of California San Diego School of Medicine, San Diego, CA, United States
[8]Department of Microbiology and Immunology, University of Nevada Reno School of Medicine, Reno, NV, United States
[9]Institute of Pathology, University Hospital Basel, Basel, Switzerland
[10]Department of Pathology, Brigham and Women's Hospital, Harvard Medical School, Boston, MA, United States
[11]Broad Institute of Harvard and MIT, Cambridge, MA, United States
[12]Department of Pathology, Dana Farber Cancer Institute, Boston, MA, United States
[+]Co-first authors
[*]Co-second authors
[†]Senior authors
[‡]Lead contacts

MANUSCRIPT

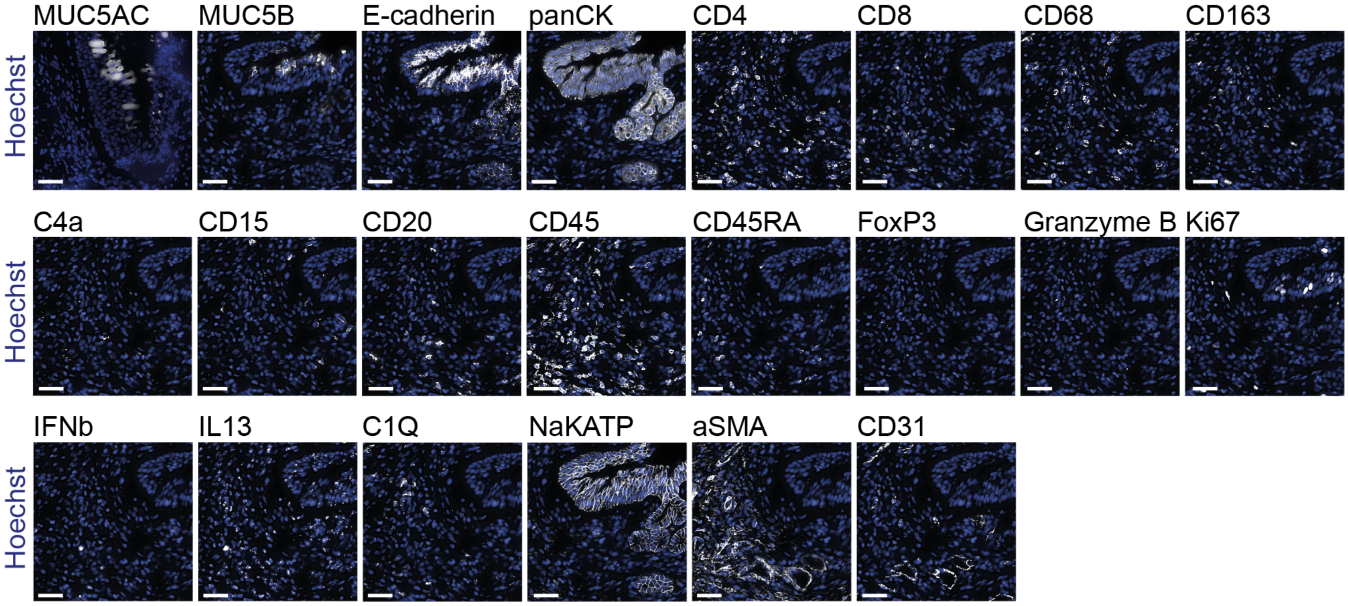


**Figure S1, related to Figure 4. Validation of CODEX antibody staining specificity.** Representative CODEX images showing 22 antibody markers (white) overlaid with the cell nucleus stain Hoechst. Scale bar: 40 µm.

### Phenotype Masks

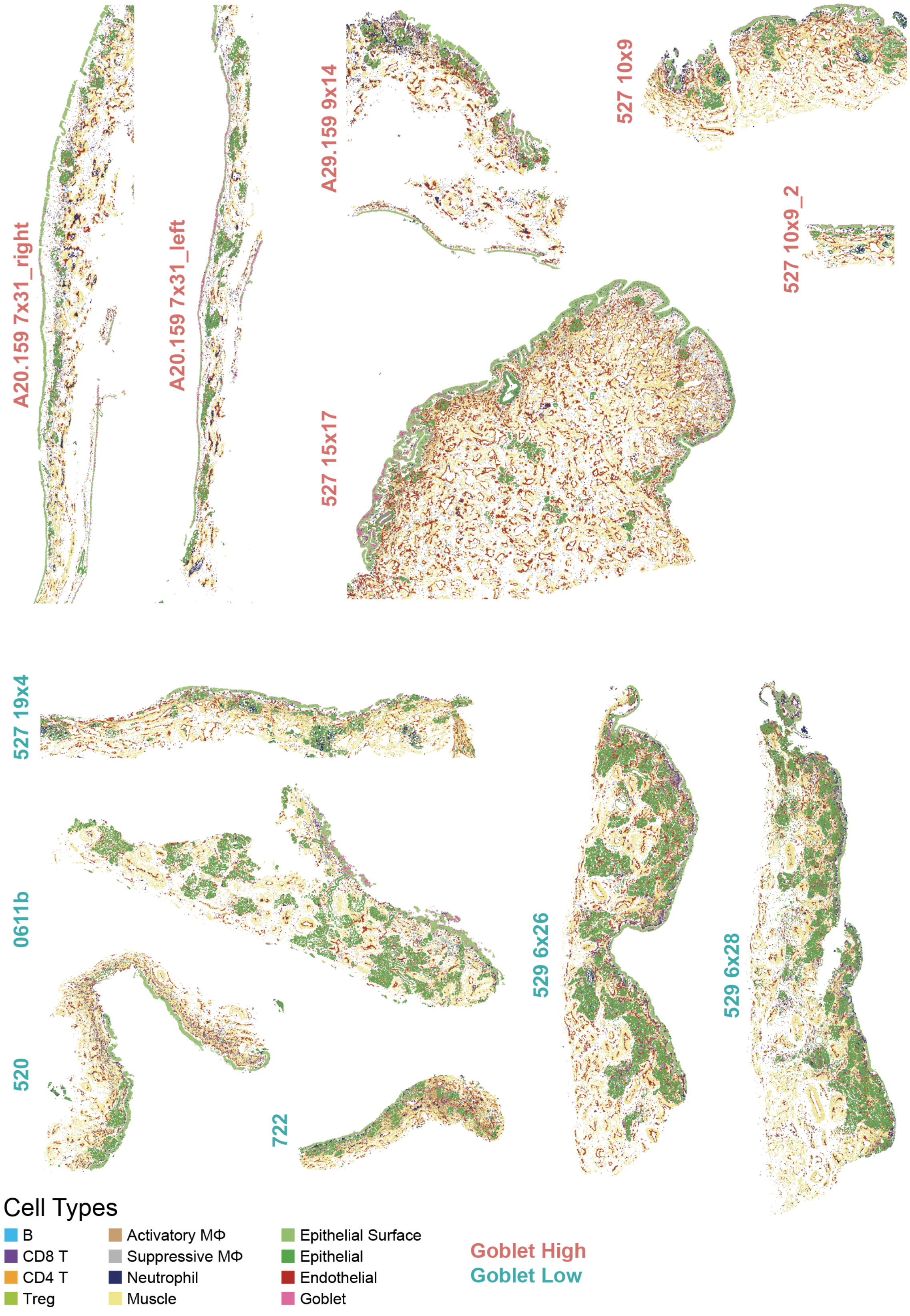


**Figure S2, related to Figure 4. Cell phenotype maps.** Phenotype maps of all nasal tissue sections.

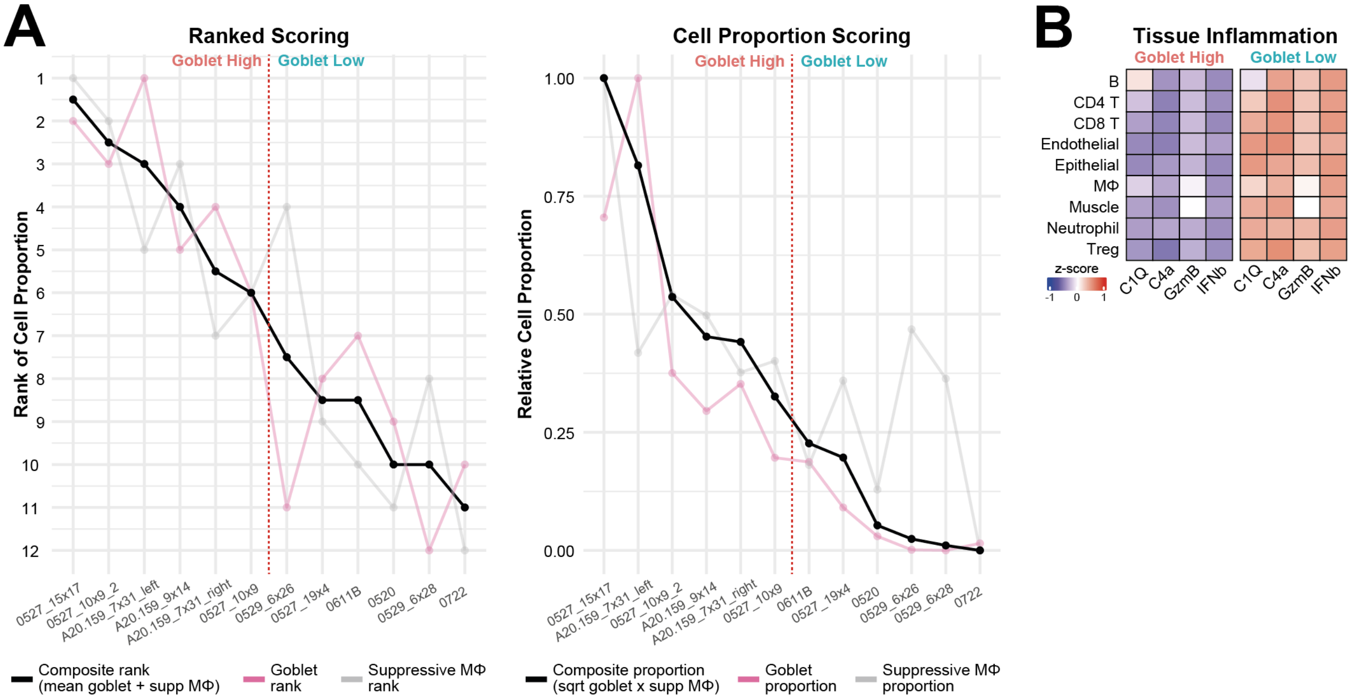


**Figure S3, related to Figures 4 & 5. Stratification into Goblet High and Goblet Low groups. (A)** Relative enrichment of goblet cells and suppressive macrophages across the sampled tissues. Left: Tissues ranked by goblet and suppressive macrophage proportion, with the composite rank calculated as the average of the two ranks. Right: Normalized goblet cell and suppressive macrophage proportions for each tissue, with the composite proportion calculated by multiplying these proportions. The red dotted lines indicate the median cutoff used to stratify tissues into high and low-enrichment groups, which returned the same results. **(B)** Heatmap of immune inflammatory marker expression across Goblet High and Goblet Low tissues.

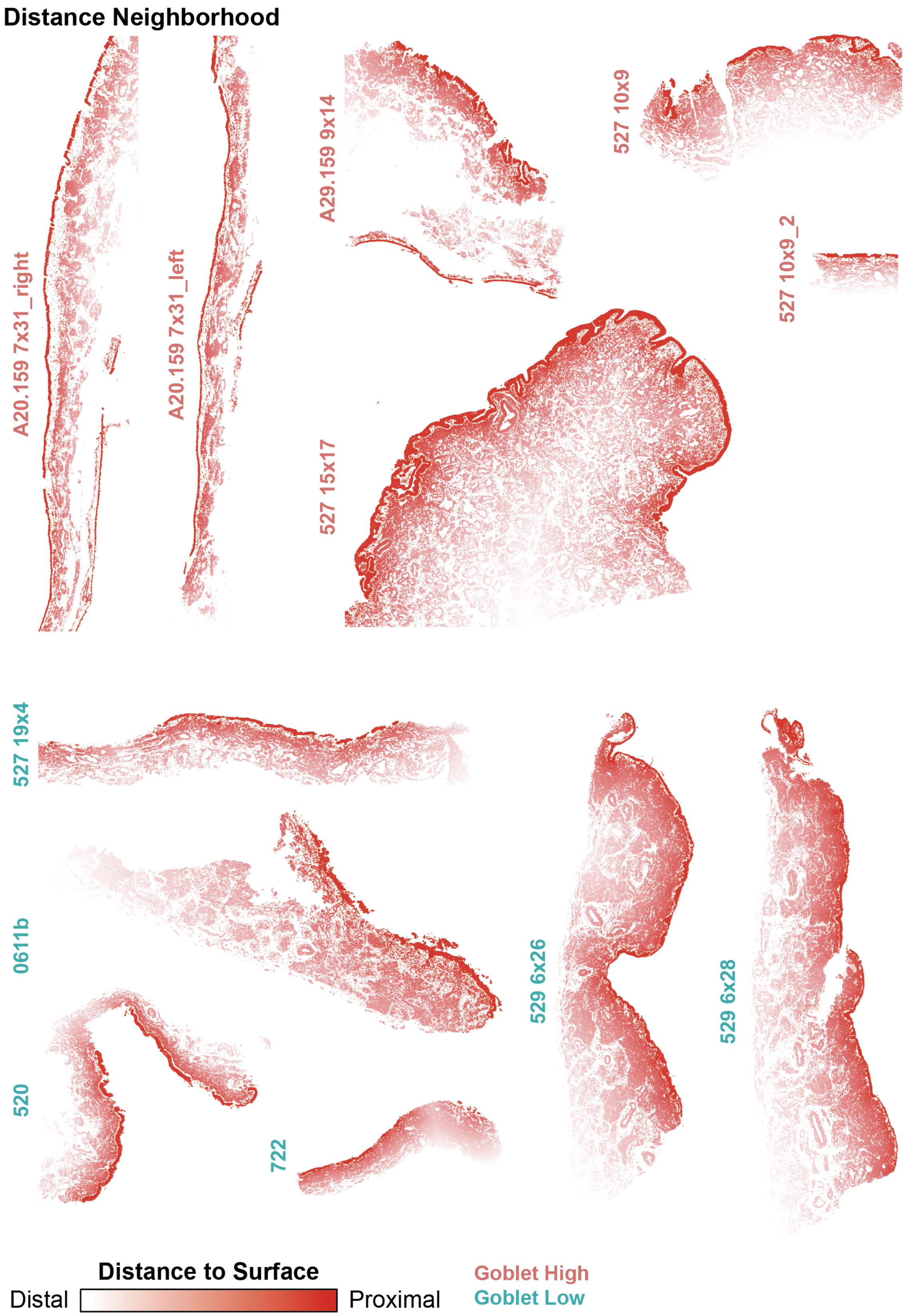


**Figure S4, related to Figure 5E. Distance neighborhood gradients.** Distance neighborhoods of all nasal tissue sections, with the surface epithelium set as the most proximal layer.

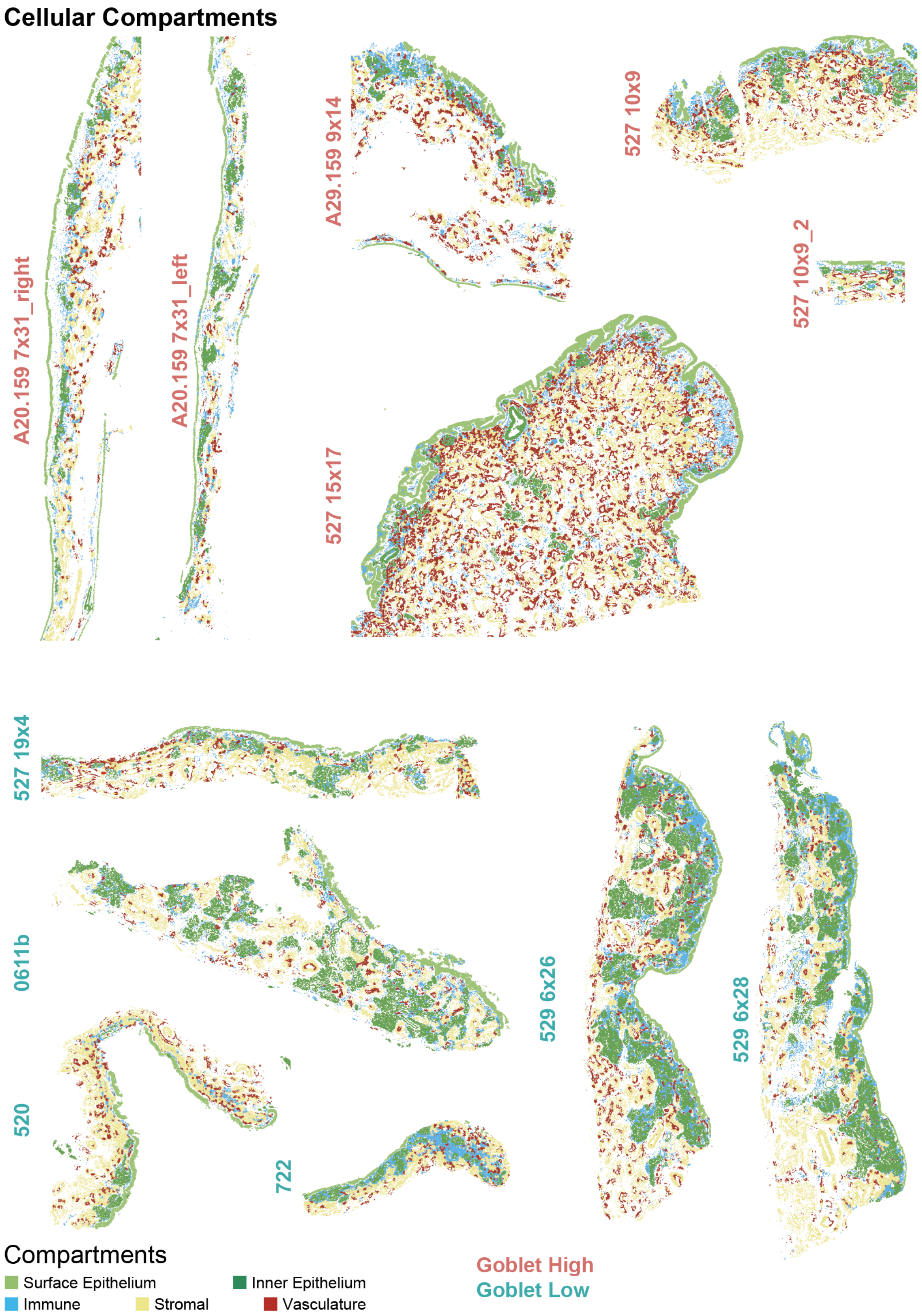


**Figure S5, related to Figure S6A. Local cell compartments.** Compartmentalization of cell phenotypes across all nasal tissues. Note that the surface epithelium compartment was set as the starting point (bin 0) and thus not seen in **Supp Fig. 5B**.

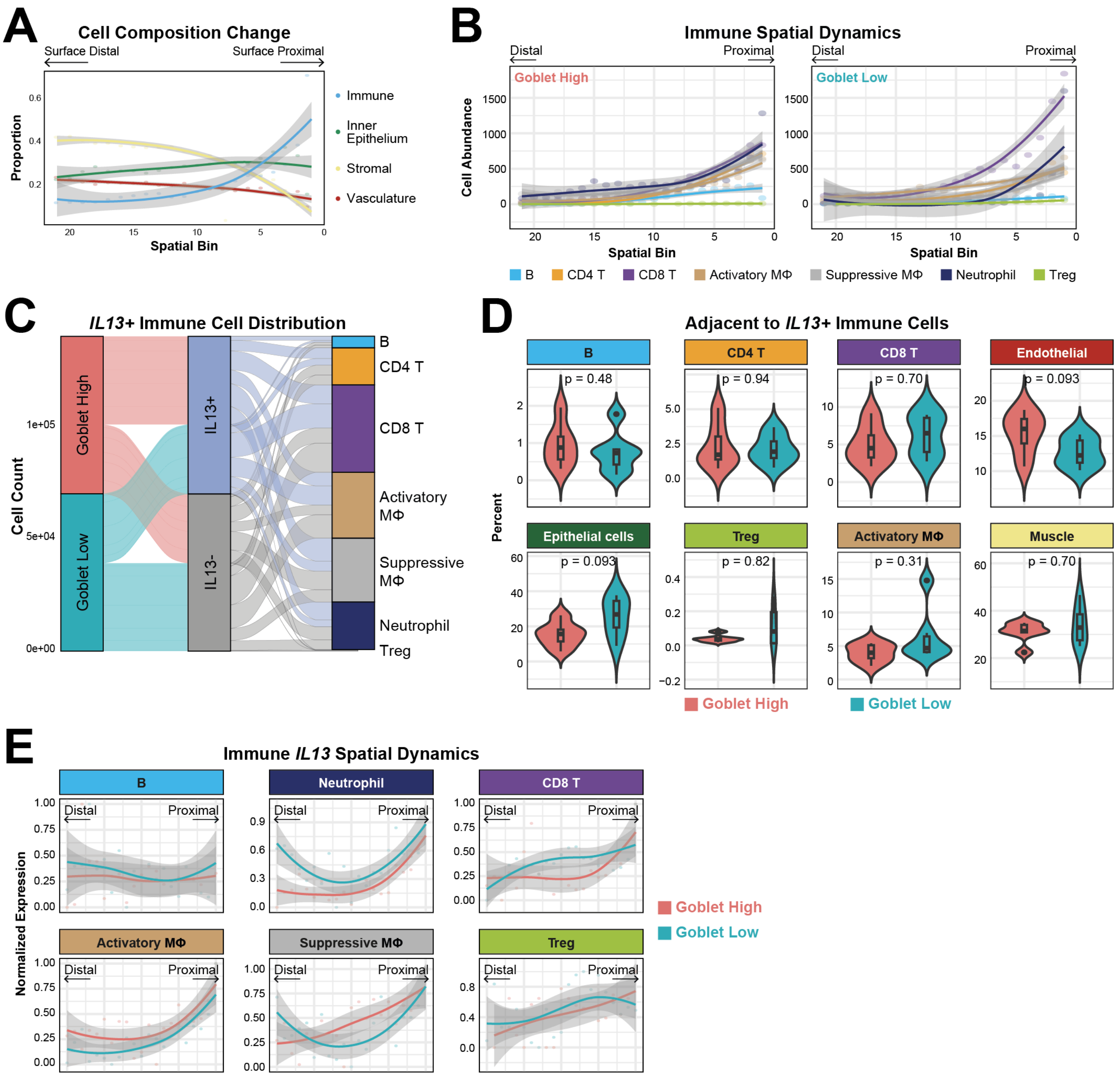


**Figure S6, related to Figure 5. Spatial differences between Goblet High and Goblet Low tissues.** **(A)** Changes in the abundance of cellular compartments **(Supp Fig. 4)** from surface-distal to surface-proximal layers. Each dot represents proportion of cells belonging to the respective cellular compartment within a given spatial bin. **(B)** Relative immune cell abundance towards the surface epithelium. Each dot represents average abundance of the respective immune cell type within a given spatial bin. **(C)** Sankey plot showing the distribution of *IL13*-positive and *IL13*- immune cells in Goblet High and Goblet Low tissues. **(D)** Abundance of cell types adjacent to *IL13*+ immune cells that do not show statistically significant differences between Goblet High and Goblet Low tissues. **(E)** Relative expression of *IL13* across other immune cells from surface-distal to surface-proximal layers between Goblet High and Goblet Low tissues. Each dot represents average *IL13* expression in the respective immune cell within a given spatial bin. Grey boundaries indicate 95% confidence interval.

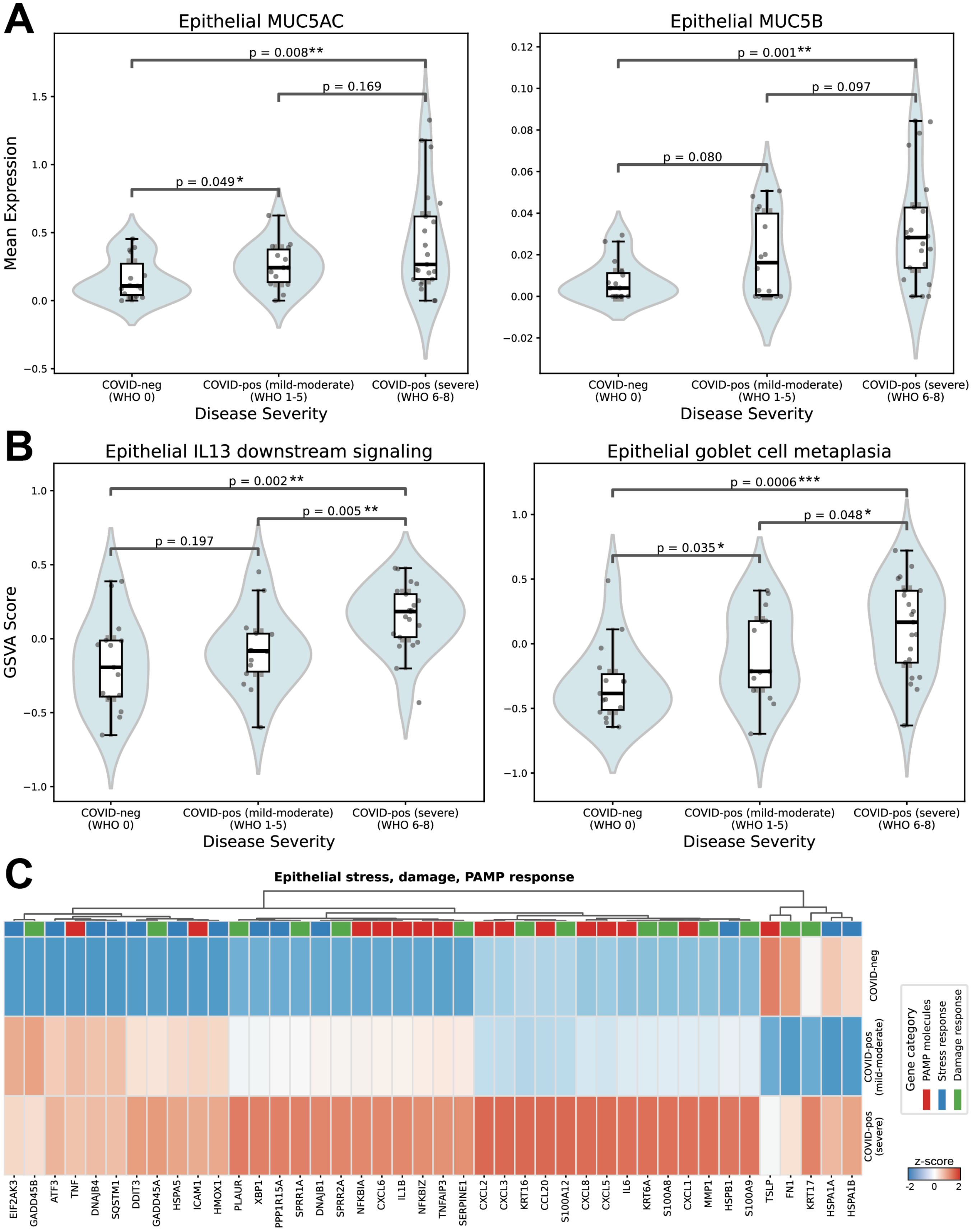


**Figure S7, related to Figure 5. COVID-19 disease severity correlates with mucin, IL13 response, and goblet cell metaplasia in the nasopharynx epithelium. (A)** *MUC5AC* (left) and *MUC5B* (right) gene expression. **(B)** GSVA score of IL13-responsive (left) and goblet cell metaplasia (right) gene signature. One-sided Wilcoxon rank-sum tests were conducted for each comparison (* $p \leq 0.05$, ** $p \leq 0.01$, *** $p \leq 0.001$), with the alternative hypothesis that that the first group has lower values than the second group. Each data point represents the mean value per patient. **(C)** Z-score expression heatmap of representative PAMP, stress, and damage response genes.

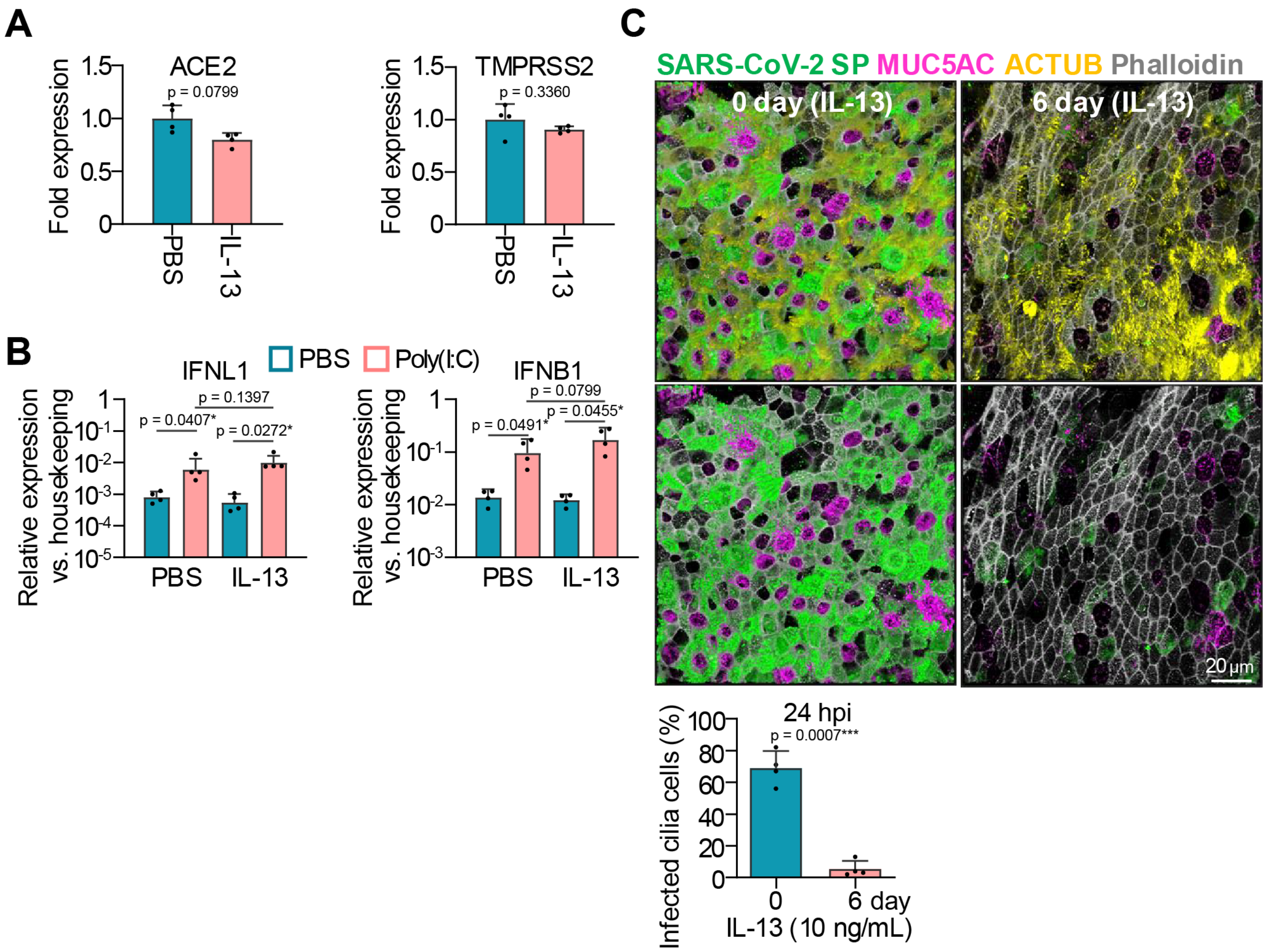


**Figure S8, related to Figure 7. IL13 has minimal effects on the expression of SARS-CoV-2 entry factors. (A)** ACE2 and TMPRSS2 mRNA expression in PBS- and IL13-treated HNEs measured by RT-qPCR. **(B)** Poly(I:C)-induced interferon responses in PBS- and IL13-treated HNEs. Cultures were treated with poly(I:C) (10µg/mL) in the basal medium for 24 h and analyzed by RT-qPCR. Data are shown as $2^{-\Delta Ct}$ values normalized to a housekeeping gene. **(C)** HNE cultures were treated with IL13 for 6 days and infected apically with SARS-CoV-2 (MOI 5, 10µL). At 18hpi, apical virus was collected in 100µL PBS, and cultures were fixed for immunofluorescence analysis. Representative IF images are shown for SARS-CoV-2-infected HNEs. Quantification of the percentage of SARS-CoV-2-positive ciliated cells is shown at bottom. Each dot represents one donor; error bars indicate mean ± SD. Statistics: paired two-tailed Student's t test (A and C) or paired one-way ANOVA with Tukey's multiple-comparisons test (B). * $p \leq 0.05$, ** $p \leq 0.01$, *** $p \leq 0.001$.
